# Investigating the evolutionary origins of the first three SARS-CoV-2 variants of concern

**DOI:** 10.1101/2022.05.09.491227

**Authors:** Mahan Ghafari, Qihan Liu, Arushi Dhillon, Aris Katzourakis, Daniel B Weissman

## Abstract

The emergence of Variants of Concern (VOCs) of SARS-CoV-2 with increased transmissibility, immune evasion properties, and virulence poses a great challenge to public health. Despite unprecedented efforts to increase genomic surveillance, fundamental facts about the evolutionary origins of VOCs remain largely unknown. One major uncertainty is whether the VOCs evolved during transmission chains of many acute infections or during long-term infections within single individuals. We test the consistency of these two possible paths with the observed dynamics, focusing on the clustered emergence of the first three VOCs, Alpha, Beta, and Gamma, in late 2020, following a period of relative evolutionary stasis. We consider a range of possible fitness landscapes, in which the VOC phenotypes could be the result of single mutations, multiple mutations that each contribute additively to increasing viral fitness, or epistatic interactions among multiple mutations that do not individually increase viral fitness—a “fitness plateau”. Our results suggest that the timing and dynamics of the VOC emergence, together with the observed number of mutations in VOC lineages, are in best agreement with the VOC phenotype requiring multiple mutations and VOCs having evolved within single individuals with long-term infections.

## Introduction

For the first 8 months of the SARS-CoV-2 pandemic, the virus exhibited a very slow pace of adaptation, with D614G being the only persistent adaptive substitution that appears to have resulted in an increased transmissibility of the virus [1–3]. However, during the second half of 2020, three designated variants of concern (VOCs) of SARS-CoV-2, Alpha, Beta, and Gamma, emerged independently and in quick succession [4–6]. No other VOC emerged until Delta and Omicron in 2021 which appear to be very different, both genetically and phenotypically, from the three original VOCs [7–8]. The VOCs are characterised by a large number of mutations relative to the genetic background from which they first emerged, and exhibit altered phenotypes resulting in varying combinations of increased transmissibility, virulence, and immune evasion [6, 9–11].

Phylogenetic analyses show that a large number of mutations, mostly located in the spike protein, have independently evolved in multiple lineages of SARS-CoV-2 including the Alpha, Beta and Gamma variants and are likely playing a key role in the adaptive evolution of the SARS-CoV-2 [7–12]. Experimental measurements and molecular dynamics simulations also show that some of these mutations have synergistic interactions for important functional traits [13–14], indicating that they may have greater combined fitness benefit to the virus. Some of the distinctive mutations in the VOCs, including the E484K and N501Y mutations found in the first three VOCs, have also been observed in chronic infections such as those in certain immunocompromised individuals [15–17], suggesting that the VOCs may have arisen from such infections. Some of the other possible explanations for the emergence of VOCs include prolonged circulation of the virus in areas of the world with poor genomic surveillance or reverse-zoonosis from other animals such as rodents followed by sustained transmission and adaptive evolution within the animal population and a spill over back to the humans (see [18] for a recent review on the possible origins of variants of SARS-CoV-2).

While finding the evolutionary process(es) that may have led to the emergence of VOCs has profound consequences for understanding the fate of the SARS-CoV-2 pandemic, there have currently been no systematic investigations to assess the likelihood of any particular evolutionary pathway that would lead to the emergence of VOCs. In this work we investigate whether the emergence of VOCs was the result of evolution via sustained transmission chains between acutely infected individuals or prolonged infections, and evaluate plausible fitness landscapes. We also discuss the potential implications of our results for the future of the pandemic and potential measures that might lower the rate at which new VOCs emerge.

## Results

### Emergence of VOCs: an evolutionary puzzle

The Alpha, Beta, and Gamma VOCs arose independently and in quick succession, with several shared mutations, in three different countries and began to spread globally (**Figure 1**). This long waiting time followed by clustered emergence of a handful of lineages was not predicted by any simple evolutionary theories. Typically, one would assume that either the beneficial mutation supply is small, in which case one expects a long waiting time for the first VOC but also long gaps before subsequent VOCs, or the mutation supply is large, in which case one expects many VOCs with only a short waiting time [19]. Moreover, each VOC had >6-10 mutations distinguishing it from then-dominant genotypes, which was also unexpected. One of the key evolutionary questions is whether VOCs evolved over the course of many acute infections or within single chronic infected hosts. Both possibilities have serious issues. The many-acute-infections hypothesis needs to explain how the virus acquired so many changes, as the mutant lineages would have had to remain at frequencies below the detection threshold in different countries for several months. The chronic-infection hypothesis needs to explain both why adaptation to the within-host environment led to a transmission advantage between hosts, and why there was no ‘leakage’ of some intermediate mutations at the between-host level before the emergence of the VOCs, i.e., why genotypes with some of the VOC mutations did not escape from the chronically infected patients earlier.

**Figure 1:**
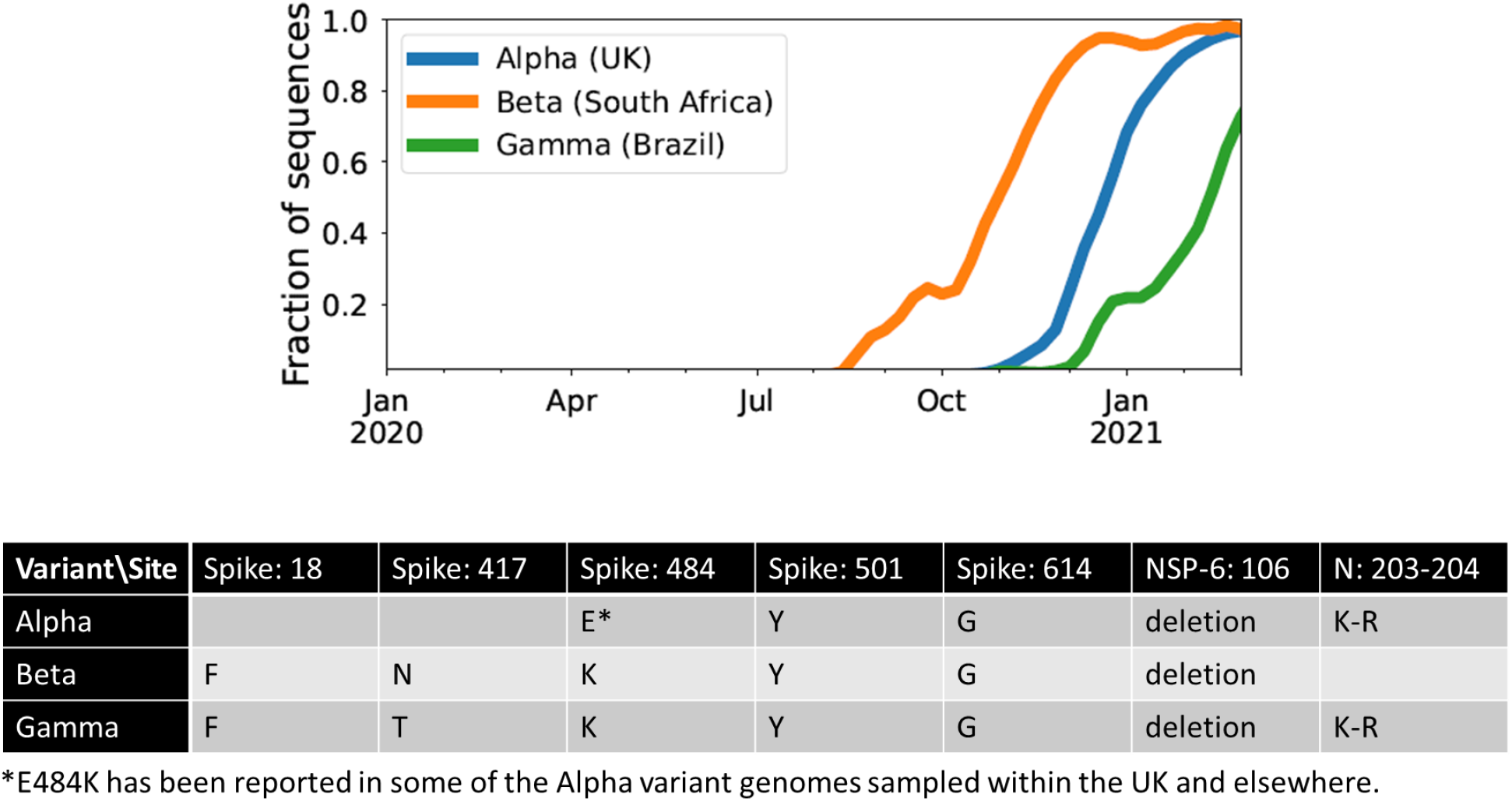
The three initial Variants of Concern arose in quick succession after a long period of limited adaptation. For each VOC, the curve shows its frequency among the SARS-CoV-2 sequences collected each week from its country of origin. The table shows the amino acid changes across the SARS-CoV-2 genome that are shared between at least two of the three VOCs [7].

### Between-host model of VOC emergence

We assume the effective virus population size is *N*_e_=*N/σ^2^* where *N* is the number of infectious individuals worldwide and *σ^2^* is the variance in offspring number (secondary cases). We treat each acute infection as one generation, assuming a tight transmission bottleneck of a single virion [20–22]. Viruses mutate at rate *μ* per base per generation (see **Methods** section). For a mutant virus population with selective advantage *s* relative to the background, the average number of secondary cases increases by a factor 1+*s*. We also assume that the number of secondary cases approximately follows a negative binomial distribution with mean *R_t_* and dispersion parameter *k*, so that *σ^2^*≈*R_t_*(1+*R_t_/k*). There is substantial uncertainty in the amount of overdispersion in the pandemic, and consequently similar uncertainty in the effective population size. Therefore, we consider a range of values for *k* to see if any would be consistent with the observed dynamics of the VOC emergence. We also note that while the importance of spatial structure is clearly visible in the spatially restricted initial spread of the VOCs from real-world data, we expect that we can neglect it when analysing their *emergence*. This is because spatial structure should not have a large impact on viral dynamics until a lineage becomes locally common, and the specific mutations differentiating the VOCs were all locally rare prior to their emergence.

### Within-host model of VOC emergence

Unlike tracking the between-host evolution of SARS-CoV-2 where an unprecedented effort has led to huge numbers of consensus genome sequences [23], our current knowledge of the within-host evolutionary dynamics of SARS-CoV-2 is still very limited, particularly in those with chronic infections. Because there is very limited data with which to constrain the within-host evolutionary dynamics of chronic infections with SARS-CoV-2, we simply treat it as a ‘black box’ and assume with some probability, *P*_f_, that a new infection is chronic and may lead to the production of a VOC (**Table 1**; **Methods** section). We also assume that within-host substitutions required for the production of the VOC occur at a constant rate *μ*_C_ per generation (see **Table 1**). (Here a generation is still defined as the typical length of an *acute* infection.) Given that we know only three VOC lineages emerged by late 2020, we expect *T*_obs_*N P*_f_~3 where *T*_obs_~180-317 days is the expected time to the emergence of the first VOC since the beginning of the pandemic based on phylogenetic estimates (see **Table 1**). Therefore, given the typical variation in the population size throughout the pandemic for biologically relevant parameter combinations *N*~1×10^6^-1×10^7^, we expect that values of *P*_f_~5×10^-9^-1×10^-7^ will maximize the likelihood of the within-host model and focus on these.

**Table 1:**
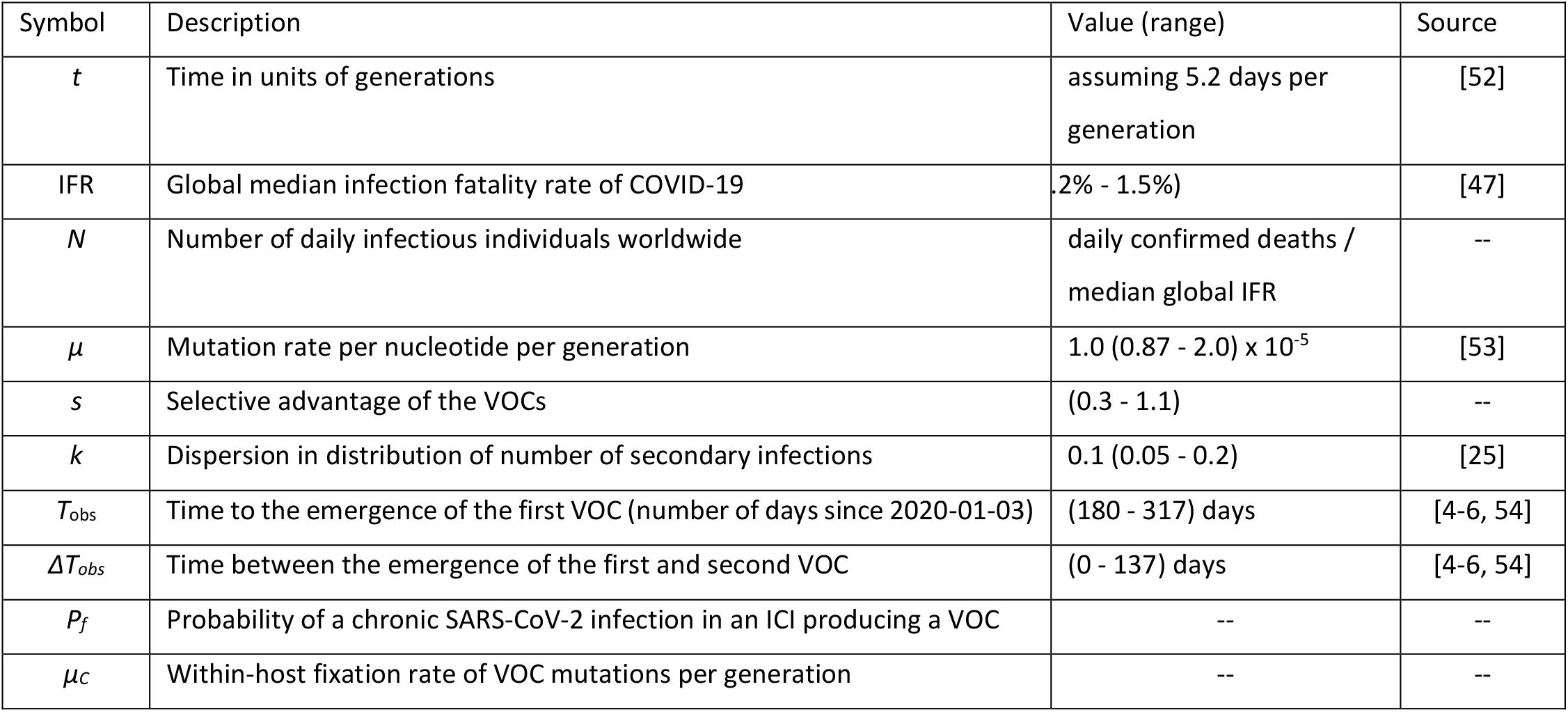
Model parameters.

### Fitness landscapes

One possible explanation for the temporal clustering of VOCs with large numbers of mutations is that the underlying fitness landscape may have some structure that causes the dynamics to deviate from our usual expectations. Unfortunately, the full space of possible fitness landscapes is enormous and impossible to explore exhaustively. To investigate the possible effects of the landscape on the dynamics, we therefore focus on three limiting local fitness landscapes that span a range of biologically plausible scenarios (**Figure 1A**). Importantly, these landscapes describe only between-host fitness, which could be very different from within-host fitness. As mentioned above, we treat within-host dynamics implicitly using an effective substitution rate and so do not need an explicit fitness landscape for it. In all three landscapes, the peak is a VOC phenotype with fitness advantage *s* over the ancestor. We assume that Alpha, Beta, and Gamma are similar enough that they can be approximately described by the same landscape and the same value of *s*, which we infer from the early rate of increase of the VOCs (see **Methods**). Landscape 1 is the simplest possibility: a single mutation on the ancestral background is sufficient to confer the full advantage. In Landscape 2, we test whether simply increasing the number of mutations involved can explain the temporal clustering. In this landscape, the VOC phenotype is produced by a combination of *K* > 1 mutations, each providing an independent fitness benefit *s/K*. In Landscape 3, we test whether epistasis may have an effect: the VOC phenotype again requires *K* mutations, but we now assume that they provide no fitness benefit until the full combination is acquired, i.e., the population must cross a fitness plateau. As mentioned above, there is experimental evidence for this form of epistasis among the VOC mutations [13–14]. We expect that shallow fitness valleys will produce similar dynamics to Landscape 3, as will shallow upward slopes with a large jump in fitness at the end [24]. Note that mutations in all the three landscapes can be acquired via the between- or within-host evolutionary pathways (**Figure 1B**). For each evolutionary scenario, we test whether there are parameter values consistent with the data on the timing of the emergence of Alpha, Beta, and Gamma variants of SARS-CoV-2 (see **Methods**; **Table 1**). For these parameter values, we further investigate whether they correspond to biologically reasonable scenarios in terms of the frequencies of the intermediate mutations prior to the emergence of VOCs, total number of mutations required to produce VOCs, total number of successful VOC lineages produced over time, and the timing between the emergence of different VOC lineages.

#### Landscape 1: single mutations

We start with the simplest possible fitness landscape, in which a single mutation conferring a fitness advantage *s* relative to the genetic background of circulating lineages is required for the emergence of VOCs. We first consider the between-host evolutionary pathway. As long as the effective population size of the pandemic was not much smaller than the census size (i.e., overdispersion was not too large), the mutation supply *N*_e_ *μ* became large early in 2020. At this point, numerous lineages would have emerged over a short period of time (see the *k*=0.2 scenario in **Figure 3A**), inconsistent with the observed dynamics. We can therefore rule out this scenario. If overdispersion were very large, it could have kept *N*e *μ* low through the establishment of the VOCs (see the *k*=0.005 and 0.001 scenarios in **Figure 3A**). **Figure 3** shows that under extremely high levels of overdispersion (*k*=0.005 and 0.001) this model can match the long waiting time for the emergence of the first VOC. However, such high levels of overdispersion are not supported by any existing epidemiological studies on SARS-CoV-2 transmission [25]. Moreover, **Figure 3B** shows that this model rarely produces an evolutionary dynamics that would fit the joint waiting time distribution for all three VOCs (also see **Supplementary Figure 1**). Under these mutation-limited conditions, there is an approximately exponential waiting time for the arrival of each VOC lineage (once we reach the point where COVID-19 becomes a pandemic in March 2020). Thus, it predicts similarly long waiting times for the emergence of Alpha, Beta, and Gamma, inconsistent with the observed temporal clustering. Therefore, there is no biologically reasonable combination of parameters that result in the clustered emergence of VOCs in late 2020 via the Landscape 1 between-host evolutionary pathway.

**Figure 2:**
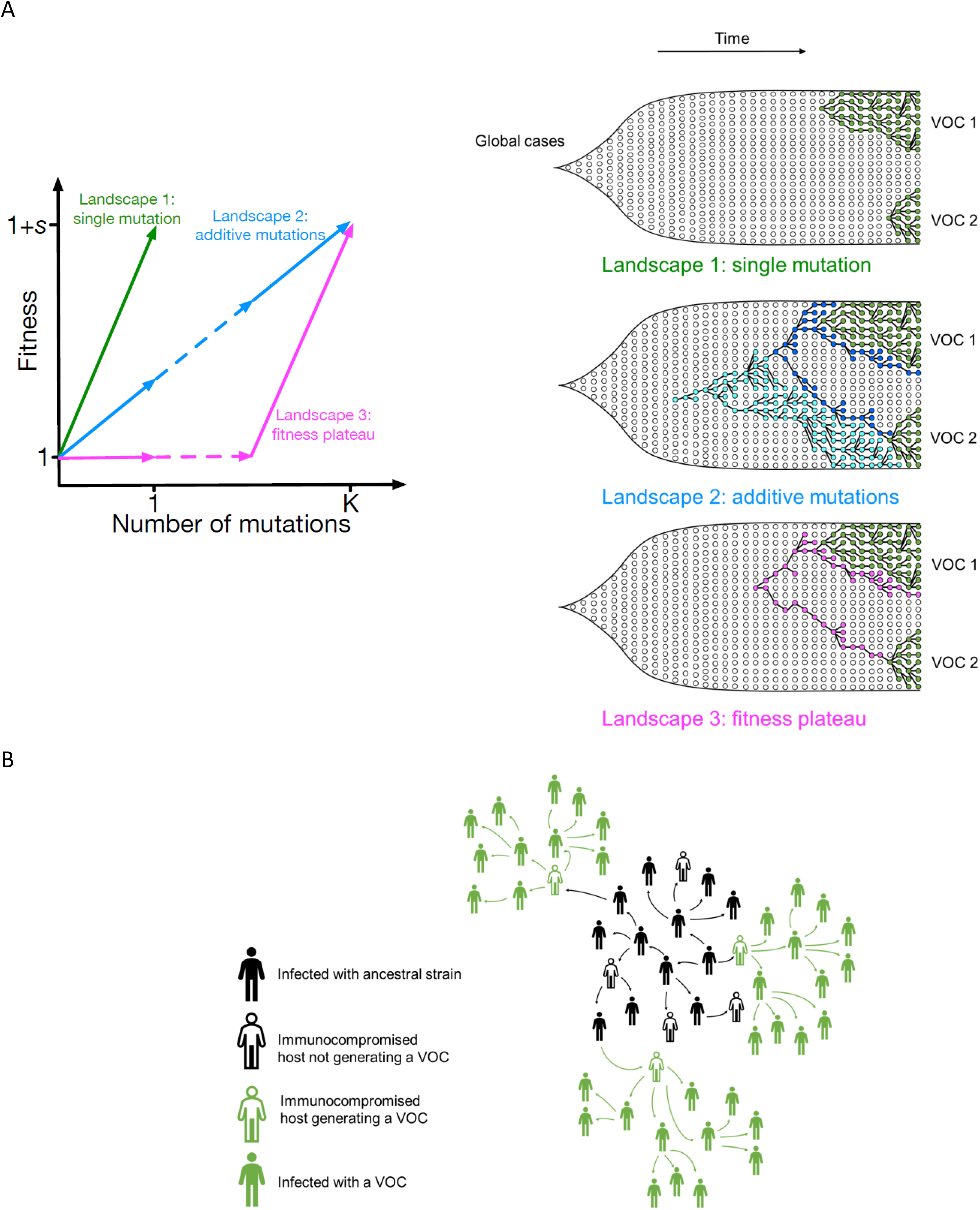
Possible evolutionary pathways to the emergence of SARS-CoV-2 VOCs. **(A**, left) The three limiting fitness landscapes for the emergence of VOCs as a function of the relevant number of mutations required, *K*. **(A**, right) VOCs can emerge from either a single advantageous mutation (green) or multiple mutations that each contribute independently to increasing fitness (blue) or only in combination (magenta). **(B)** Emergence of VOCs via the within-host evolutionary path such that an infectious individual passes on a wild-type variant of the virus to an immunocompromised individual where the virus may acquire the relevant mutations during the chronic phase of the infection and later be passed on to the rest of the population.

**Figure 3:**
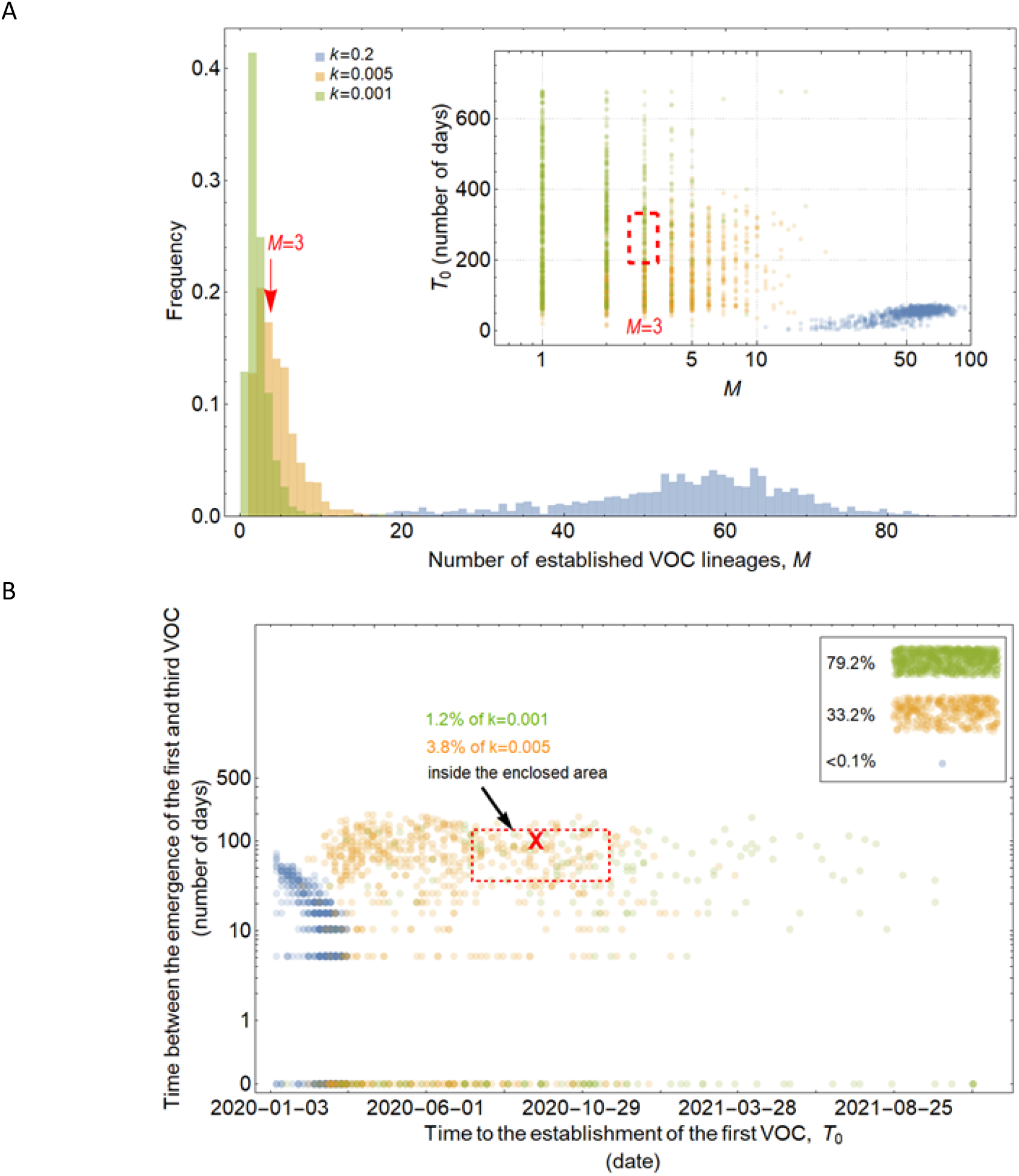
Evolution between hosts on a single-mutation landscape (*K* = 1) rarely reproduces the observed VOC dynamics, even with extreme overdispersion. **(A)** Total number of established VOC lineages (*M*) measured under varied levels of overdispersion, *k*, such that IFR=1.5%, *μ*=0.87×10^-5^, *K*=1, and *s*=0.4. The inset shows *M* with respect to the waiting time for the establishment of the first VOC lineage since the start of the pandemic, *T*_0_. The region corresponding to the waiting time for the emergence of the first three SARS-CoV-2 VOC is highlighted in red. Under low levels of overdispersion (blue, *k*=0.2), too many VOC lineages are produced very early on in the pandemic. On the other hand, as we increase overdispersion (orange and green), fewer VOCs can establish in the population. It also takes them much longer to establish and reach high frequencies in the population. **(B)** Evaluating the temporal clustering of the first three VOC lineages. For each simulation run, represented by a point on the graph, we measure *T*_0_ and the time difference between the establishment of the first and third successful VOC lineages. The red dashed rectangle shows the region corresponding to the emergence of the first three SARS-CoV-2 VOCs with the cross sign (“**X**”) representing the mean value. We see that as the level of overdispersion increases, the emergence time of VOCs are more scattered and rarely exhibit temporal clustering in late 2020 -- Only 1.2% and 3.8% of the evolutionary dynamics corresponding to overdispersion *k*=0.005 and *k*=0.001 fall inside the enclosed area, respectively. The inset shows that 33.2% and 79.2% of the runs for *k*=0.005 and *k*=0.001 scenarios produce fewer than three successful VOC lineages by the end of the simulation period. Each run stops once the frequency of the VOC population reaches 75%. See also supplementary figure 1.

On the other hand, if VOCs arose from chronic infections, then their emergence was a two-step process: first, chronic infections had to occur, and then the VOC mutation had to arise in them. The waiting time for the first step is determined by *NP*_f_, note that the number of chronic infections depends on the census size *N* rather than *N*_e_, i.e., it is insensitive to the amount of overdispersion. The second step follows an exponential distribution within each chronic host, with rate *μ*_C_. The third step, the spread of the VOC from the original chronic host to the rest of the population, then takes much less time than the first two. **Figure 4** shows that to match observed VOC dynamics we must assume that the level of overdispersion is very high (i.e., very low mutation supply, *N*_e_*μ*), effectively blocking the between-host evolutionary pathway, while simultaneously assuming that chronic infections are very frequently produced in the population (i.e., *NP*_f_~1) and that there is a relatively long waiting time before the production of each VOC mutation (*μ*_C_~0.01). However, like the between-host pathway, this scenario requires very high levels of overdispersion which makes the Landscape 1 within-host evolutionary pathway also an unlikely explanation for the emergence of VOCs (see **Supplementary Figure 2**).

**Figure 4:**
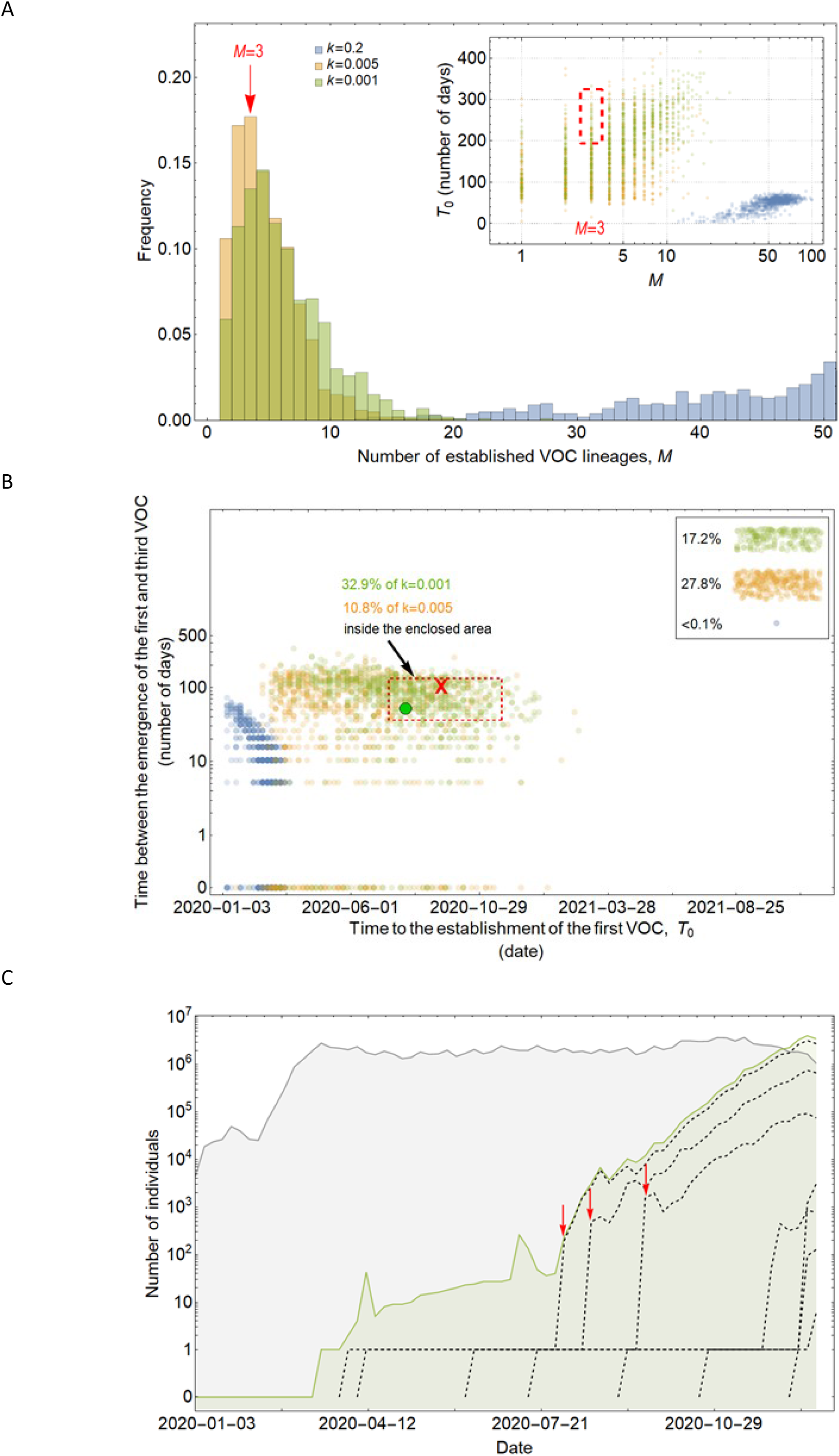
Evolution within hosts on a single-mutation (*K* = 1) landscape can match the observed VOC dynamics, but only with extreme overdispersion to prevent between-host evolution. **(A)** Total number of established VOC lineages (*M*) measured under varied levels of overdispersion, *k*, where the within-host parameters for the *k*=0.2 scenario (blue) are *P*_f_=5×10^-10^ and *μ*_C_=0.001. For the *k*=0.005 scenario (orange), *P*_f_=6×10^-8^ and *μ*_C_=0.1. Finally, for the *k*=0.001 scenario (green), *P*_f_=6×10^-6^ and *μ*_C_=0.01. For all the three scenarios, the between-host parameters *μ*=0.87×10^-5^, IFR=1.5%, *K*=1, and *s*=0.4 are the same. The inset shows *M* with respect to the waiting time for the establishment of the first VOC lineage since the start of the pandemic, *T*_0_. The region corresponding to the waiting time for the emergence of the first three SARS-CoV-2 VOC is highlighted in red. We see that under low levels of overdispersion, *k*=0.2 (blue), too many VOC lineages are produced very early on in the pandemic. On the other hand, as we increase overdispersion (orange and green), fewer VOC lineages can establish in the population, and it generally takes longer for them to do so. **(B)** Evaluating the temporal clustering of the first three VOC lineages. For each simulation run, represented by a point on the graph, we measure the time that it takes for a single adaptive mutation to establish in the population and the time difference between the establishment of the first and third successful VOC lineage. The red dashed rectangle shows the region of the parameter space corresponding to the emergence of the first three SARS-CoV-2 VOCs with the cross sign (“**X**”) representing the mean value. The graph shows that by increasing the level of overdispersion and lowering the evolutionary contribution from the between-host pathway, multiple VOCs can emerge in quick succession via the within-host pathway such that a larger fraction of the simulation runs yield the correct timing for the emergence of the first three VOCs in late 2020 (i.e., they fall inside the enclosed area). The inset shows that 27.8% and 17.2% of the runs for *k*=0.005 and *k*=0.001 scenarios produce fewer than three successful VOC lineages by the end of the simulation period. Each run stops once the frequency of the VOC population reaches 75%. **(C)** A typical evolutionary trajectory corresponding to the *k*=0.001 scenario (green) highlighted with a bold green circle in panel (B). The graph shows the VOC population (green) along with the individual VOC lineages (black dashed lines) emerging from the background population (gray). Red vertical arrows show the establishment time of the first three VOCs. We see that the VOC mutation is first produced in a single individual within the population (chronically infected case) for a relatively long time before successfully spreading to the rest of the population. See also supplementary figure 2.

#### Landscape 2: additive mutations

Landscape 2 corresponds to an evolutionary pathway in which there were *K>1* major mutations involved in the emergence of VOCs, each making an additive contribution of ≈*s/K* to fitness. If evolution occurred at the whole-population level, **Figure 5** and **Supplementary Figure 3** show that, for a range of parameter combinations, the additive fitness landscape requiring up to four mutations can create evolutionary dynamics with appropriately long waiting times before the arrival of the first successful VOC lineage, while for combinations of more than 4 mutations, VOC lineages do not emerge by late 2020 under any biologically reasonable parameter combinations for effective population size, mutation rate, and selective coefficient. However, while *K* <= 4 can match the observed waiting for the first VOC lineage, for the *K*=2 and 3, this first VOC is usually followed by the establishment of nearly a dozen VOC lineages that emerge in quick succession (see *K*=2 and 3 scenarios in **Figure 5A**; also see **Supplementary Figure 3**), inconsistent with the observation of only 3 VOC lineages emerging in late 2020. However, while for *K*=4 fewer VOC lineages are produced, a closer examination of a typical evolutionary trajectory that matches the long waiting time before the establishment of the first VOC further reveals that the intermediate single-, double-, or triple-mutants reach high frequencies before the emergence of the first successful (quadruple*-*mutant) VOC lineage (**Figure 5C**). The sequential fixation of adaptive mutations at the population level would imply that the intermediate mutations were detectable many months prior to the emergence of VOCs, again inconsistent with the genomic surveillance data from around the world. The inconsistency is also visible phylogenetically. The sequential fixation dynamics predicted by the model create a ladder-like phylogenetic relationship between the background and mutant populations whereby every new VOC mutation becomes dominant in the population before giving rise to lineages with additional mutations. Even though such phylogenetic relationships may emerge in SARS-CoV-2 over longer evolutionary timescales (as have been observed in human coronaviruses [26]), they do not resemble the observed topology of the phylogeny of the VOCs of SARS-CoV-2, which is more star-like.

**Figure 5:**
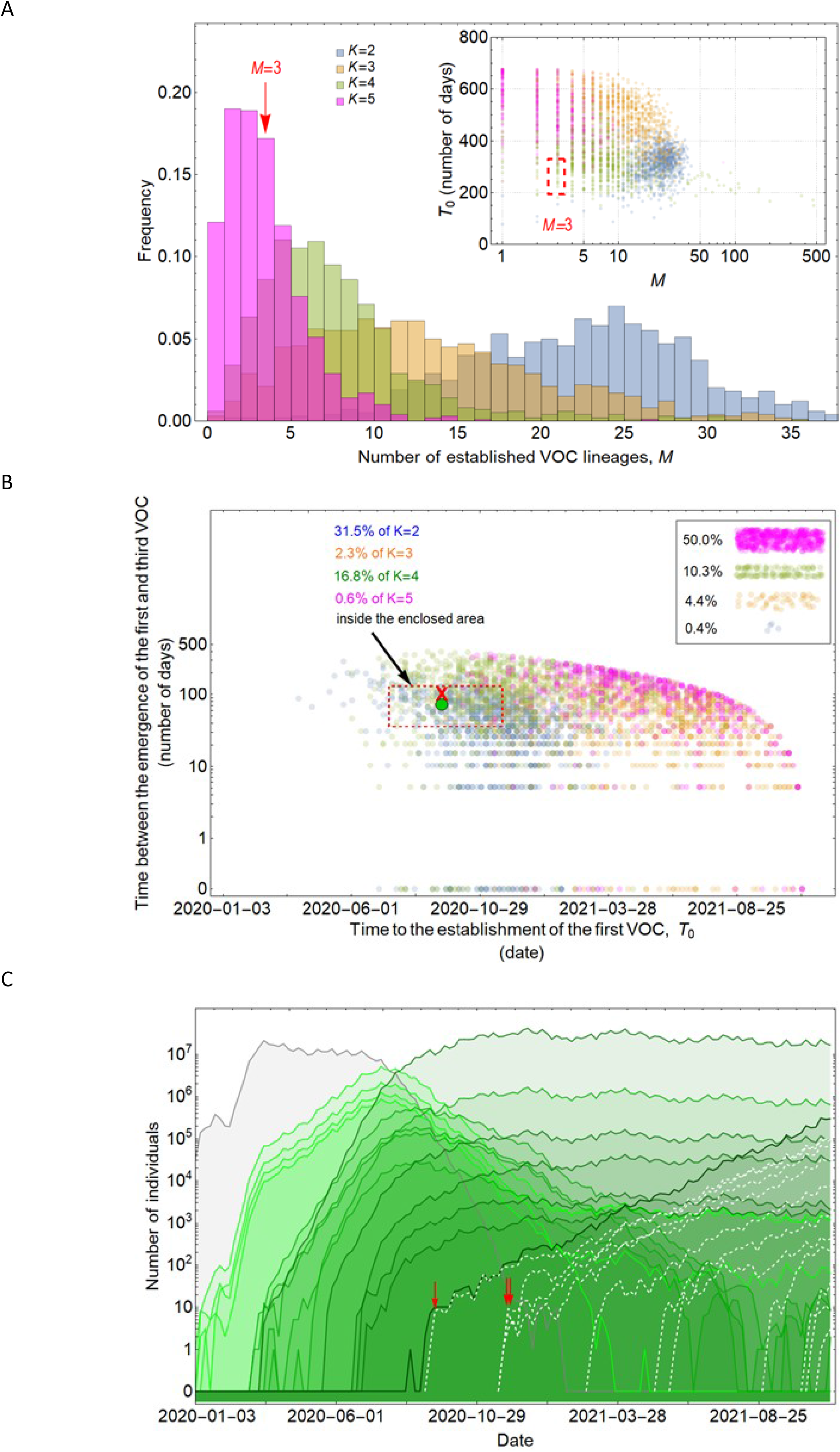
Between-host evolution on an additive fitness landscape can match the observed VOC dynamics, but only by having intermediate mutants reach unrealistically high frequencies. **(A)** Total number of established VOC lineages (*M*) for different number of mutations, *K*, involved in the production of a VOC. For the *K*=2 scenario (blue), IFR=1.5%, *μ*=0.87×10^-5^, *k*=0.05, and *s*=0.3. For the *K*=3 scenario (orange), IFR=0.5%, *μ*=0.87×10^-5^, *k*=0.05, and *s*=0.5. For both the *K*=4 (green) and *K*=4 (magenta) scenarios, IFR=0.2%, *μ*=2×10^-5^, *k*=0.2, and *s*=1.0. The inset shows *M* with respect to the waiting time for the establishment of the first VOC lineage since the start of the pandemic, *T*_0_. Under *K*=2 and 3, a very large number of successful VOC lineages are produced by late 2020, with the *K*=2 scenario producing, on average, more than 20 VOC lineages that establish in the population. On the other hand, for the *K*=5 scenario, on average, fewer than 3 lineages are produced. It also takes much longer for them to establish in the population. **(B)** Evaluating the temporal clustering of the first three VOC lineages. For each simulation run, represented by a point on the graph, we measure the time that it takes for a single adaptive mutation to establish in the population and the time difference between the establishment of the first and third successful VOC lineage. The red dashed rectangle shows the region of the parameter space corresponding to the emergence of the first three SARS-CoV-2 VOCs with the cross sign (“**X**”) representing the mean value. We see a noticeable overlap between the *K*=2 and 4 scenarios and the red rectangle suggesting that a larger fraction of the simulation runs exhibit temporal clustering dynamics for VOC emergence. The inset shows that 10.3% of the runs for the *K*=4 scenario produce fewer than three successful VOC lineages by the end of the simulation period. Each run stops once the frequency of the VOC population reaches 75%. **(C)** A typical evolutionary trajectory corresponding to the *K*=4 scenario highlighted with a bold green circle in panel (B). The graph shows the background population in gray and the *i*-mutant populations (1<*i*≤*K*) in different shades of green from light (fewer mutations) to dark (more mutations). Note that for the *K*=4 scenario, there are four single-mutant, six double-mutant, four triple-mutant, and one quadruple-mutant genotypes. The dashed lines show the dynamics of all the established VOC lineages over time. Red vertical arrows show the establishment time of the first three VOCs. We can see that some of the intermediate mutant genotypes reach close to fixation before giving rise to the VOC population. See also supplementary figure 3.

For the chronic-infection pathway, on the other hand, the intermediate mutants could have fixed within the host while remaining at undetectable frequencies at the between-host level until the production of the VOCs. **Figure 6** shows that for a combination of parameters requiring *K*=3 and 6 mutations where the mutation supply is low and the strength of selection is relatively weak such that the intermediate mutants cannot reach fixation before the emergence of the VOC population, the Landscape 2 within-host pathway can lead to the clustered emergence of a few VOC lineages by late 2020. However, if the selective coefficient *s/K* on single mutants is too high, they will reach observable frequencies before the VOCs emerge, as we discussed above with the between-host pathway. Effectively, this means that there is a minimal *K* of at least 3 needed so that the strength of selection on each mutant allele is not too strong. Alternatively, lower *K* is possible but requires extremely large overdispersion, as in the *K* = 1 case.

**Figure 6:**
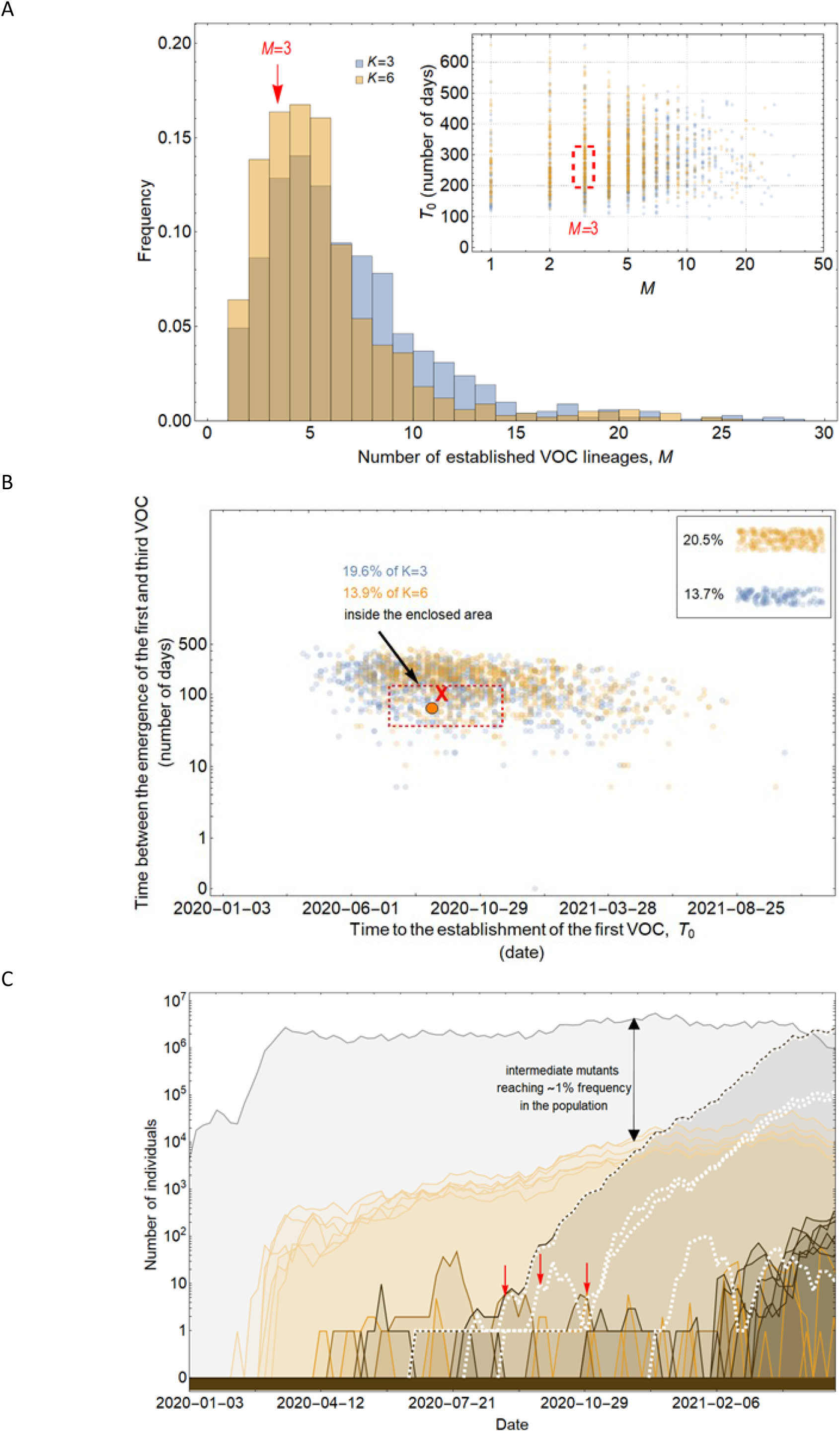
Evolution within hosts on an additive fitness landscape can match the observed VOC dynamics as long as *K* is large enough that between-host evolution is ineffective. **(A)** Total number of established VOC lineages (*M* for different number of mutations, *K*, involved in the production of a VOC. For *K*=3 (blue), the within-host parameters are *P*_f_=3.5×10^-8^, and *μ*_C_=0.15. For *K*=6 (orange), *P*_f_=3×10^-8^, and *μ*_C_=0.3. In both scenarios, the between-host parameters *μ*=0.87×10^-5^, IFR=1.5%, *k*=0.05, and *s*=0.3 are the same. The inset shows *M* with respect to the waiting time for the establishment of the first VOC lineage since the start of the pandemic, *T*_0_. The region corresponding to the waiting time for the emergence of the first three SARS-CoV-2 VOC is highlighted in red. Both scenarios produce roughly the same of number of VOC lineages. However, on average, *T*_0_ is slightly longer for the *K*=6 scenario. **(B)** Evaluating the temporal clustering of the first three VOC lineages. For each simulation run, represented by a point on the graph, we measure the time that it takes for a single adaptive mutation to establish in the population and the time difference between the establishment of the first and third successful VOC lineage. The red dashed rectangle shows the region of the parameter space corresponding to the emergence of the first three SARS-CoV-2 VOCs with the cross sign (“**X**”) representing the mean value. We can see that by having a combination of relatively high level of overdispersion, high IFR, and low between-host mutation rate, there is a lower chance of intermediate mutations reaching fixation via the between-host path. Instead, multiple VOCs can emerge in quick succession during chronic infections such that a relatively large fraction of the simulation runs yield a temporal clustering that matches the emergence of the first three VOCs in late 2020 (i.e., they fall inside the enclosed area). The inset shows that 20.5% and 13.7% of the runs for *K*=3 and 6 scenarios produce fewer than three successful VOC lineages by the end of the simulation period, respectively. Each run stops once the frequency of the VOC population reaches 75%. **(C)** A typical evolutionary trajectory corresponding to the *K*=6 scenario highlighted with a bold orange circle in panel (B). The graph shows the background population in gray and the *i*-mutant populations (1<*i*≤*K*) in different shades of green from light (fewer mutations) to dark (more mutations). The dashed lines show the dynamics of all the established VOC lineages over time. Red vertical arrows show the establishment time of the first three VOCs. We can see that the single-mutant genotypes (lines in light orange) are produced via the between-host pathway but never reach above 1% prevalence before the emergence of the VOCs (white dashed lines). See also supplementary figure 4.

#### Landscape 3: fitness plateau crossing

As in Landscape 2, Landscape 3 describes an evolutionary pathway where there are *K>1* major mutations involved in the generation of VOCs, but in this case, only the full *K*-mutant VOC genotype has a substantial selective advantage relative to the background population, while the selective advantages of the intermediate genotypes are negligible. This does not necessarily imply that the selective coefficients of the intermediate genotypes are small in the standard weak-selection sense (small relative to 1/*N_e_*), but only that they are too small to substantially affect the dynamics of the production of the first successful *K*-mutant VOC lineage, a weaker condition that depends on the mutation rate [24].

For the between-host model of VOC emergence, our analysis suggests that only a plateau-crossing of size *K*=2 may be consistent with the timing of the emergence of SARS-CoV-2 VOCs (**Figure 7**; **Supplementary Figure 5**). Extended plateaus requiring *K*>2 mutations take much longer to cross and for most parameter combinations either zero or one VOC lineage is produced before the end of 2020 (**Figure 7A**). For a typical *K*=2 plateau-crossing trajectory, single-mutant genotypes grow linearly over time and reach a frequency of ⪅0.1% before producing ~1-5 successful VOC lineages that emerge in quick succession (**Figure 7C**). Therefore, unlike the between-host evolutionary pathway in Landscape 2, a fitness plateau could have led to the clustered emergence of several VOCs after a long waiting time during which none of the intermediate mutations reached high frequency. However, the fact that for biologically plausible parameter values only a narrow plateau of *K*=2 mutations can be crossed seems inconsistent with the high number of mutations found in the VOCs and particularly with the high number of similar mutations shared across unrelated VOC lineages. This inconsistency may be partly reconciled with the possibility of compounded evolutionary effects following the plateau-crossing event such as the emergence of hyper-mutability traits across certain sites or strong within-host selection following the acquisition of the *K* mutations.

**Figure 7:**
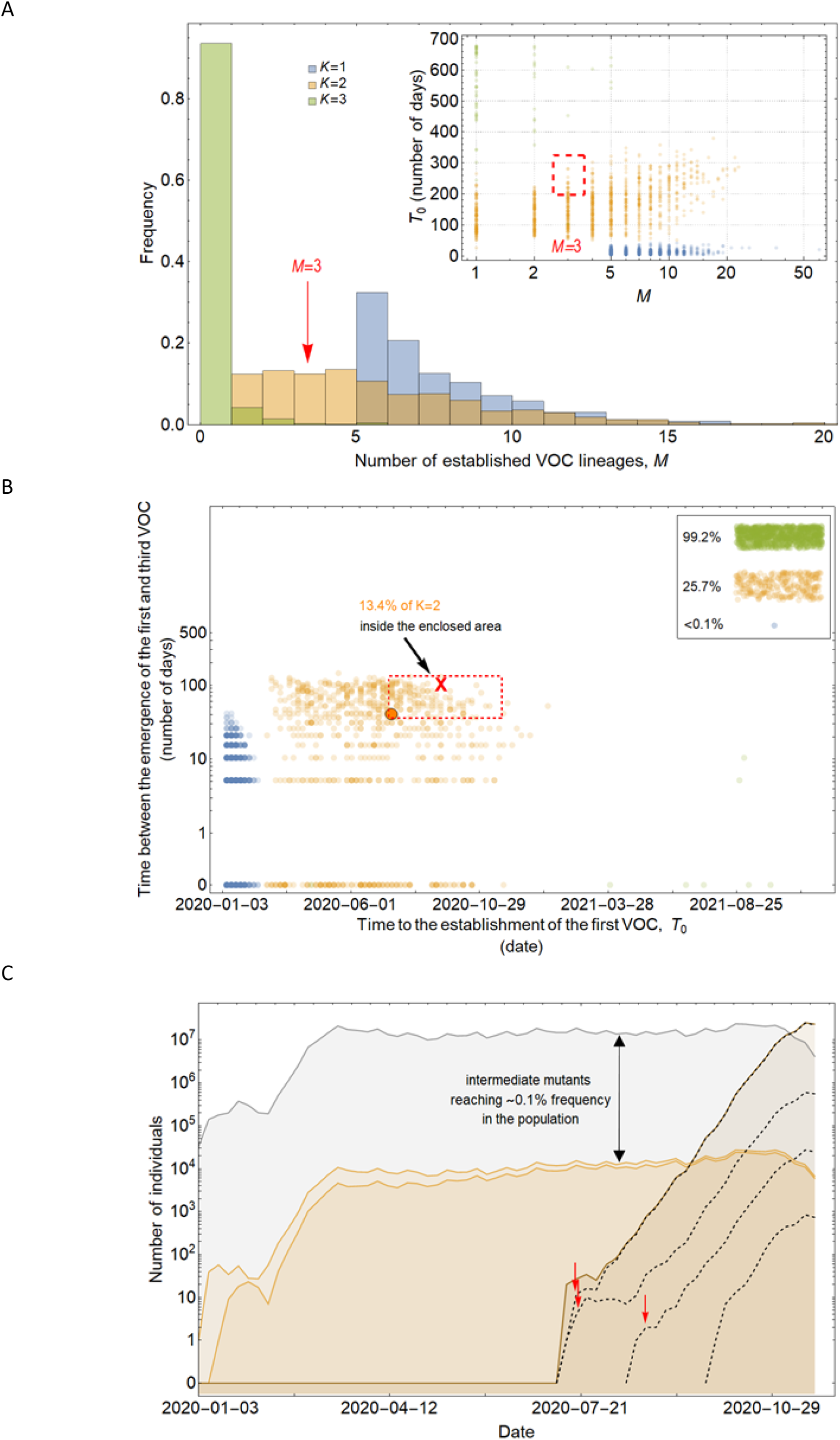
Between-host evolution on a fitness plateau can match the observed VOC dynamics, but only for *K* = 2. **(A)** Total number of established VOC lineages (*M*) for different number of mutations, *K*, involved in the production of a VOC, such that IFR=0.2%, *μ*=2×10^-5^, *k*=0.2, and *s*=1.0. The inset shows *M* with respect to the waiting time for the establishment of the first VOC lineage since the start of the pandemic, *T*_0_. For *K*=1 and 3 scenarios, there are too many and too few VOC lineages are produced by late 2020. Only for the *K*=2 scenario we can see an intermediate number of VOC lineages being produced in the right time span. **(B)** Evaluating the temporal clustering of the first three VOC lineages. For each simulation run, represented by a point on the graph, we measure the time that it takes for a single adaptive mutation to establish in the population and the time difference between the establishment of the first and third successful VOC lineage. The red dashed rectangle shows the region of the parameter space corresponding to the emergence of the first three SARS-CoV-2 VOCs with the cross sign (“**X**”) representing the mean value. We see a noticeable overlap between the *K*=2 scenario and the red rectangle suggesting that a fraction of the simulation runs exhibit temporal clustering dynamics for VOC emergence. The inset shows that 99.2% and 25.7% of the runs for the *K*=3 and 2 scenarios produce fewer than three successful VOC lineages by the end of the simulation period. Each run stops once the frequency of the VOC population reaches 75%. **(C)** A typical evolutionary trajectory corresponding to the *K*=6 scenario highlighted with a bold orange circle in panel (B). The graph shows the background population in gray, single-mutants in light orange, and double-mutants in dark orange. Note that for the *K*=2 scenario, there are two single-mutant and one double-mutant genotypes. The dashed lines show the dynamics of all the established VOC lineages over time. Red vertical arrows show the establishment time of the first three VOCs. We can see that the single-mutant genotypes reach close to 0.1% before giving rise to the VOC population. See also supplementary figure 5.

If the VOCs arose from chronic infections, the intermediate VOC mutations (which are neutral at the between-host level of selection, but may be selected within-host) can rapidly fix within a host, allowing much wider plateaus to be crossed compared to the between-host evolutionary pathway. Unlike Landscape 2 within-host pathway, the early leakage of intermediate mutations to the population is much less likely as they have no strong selective advantage over the background population. **Figure 8** shows that the within-host evolutionary pathway of Landscape 3 creates evolutionary trajectories that are consistent with the clustered emergence of ~3 VOCs in late 2020. There is also less seeding of new chronic infections with intermediate mutations, leading to fewer VOC lineages compared to Landscape 2 (also see **Supplementary Figure 6**).

**Figure 8:**
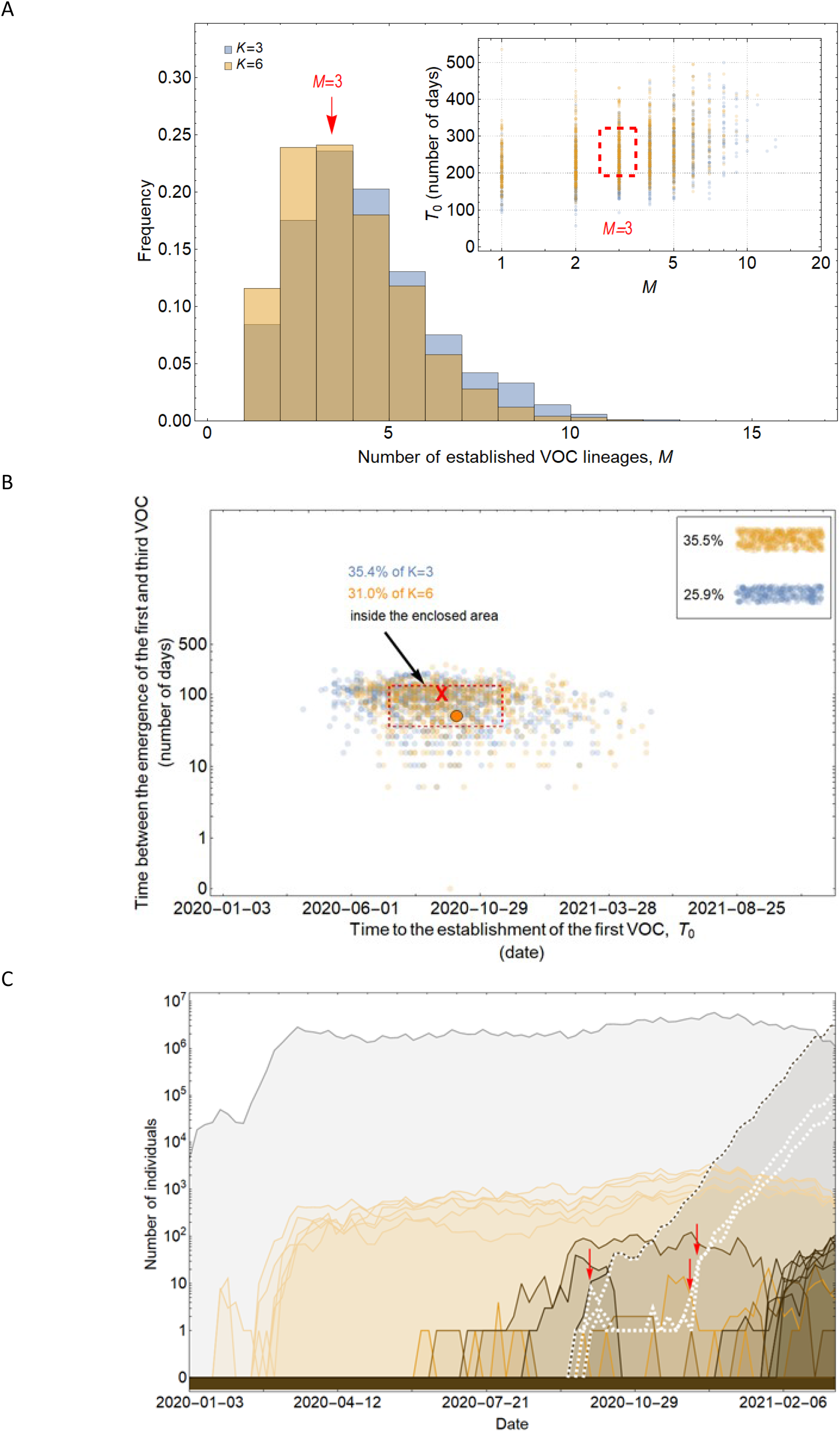
Within-host evolution on a fitness plateau can match the observed VOC dynamics for a large range of plateau widths. **(A)** Total number of established VOC lineages (*M* for different number of mutations, *K*, involved in the production of a VOC. For *K*=3 (blue), the within-host parameters are *P*_f_=2×10^-8^, and *μ*_C_=0.1. For *K*=6 (orange), *P*_f_=4.5×10^-8^, and *μ*_C_=0.25. In both scenarios, the between-host parameters *μ*=1×10^-5^, IFR=0.5%, *k*=0.1, and *s*=0.7 are the same. The inset shows *M* with respect to the waiting time for the establishment of the first VOC lineage since the start of the pandemic, *T*_0_. The region corresponding to the waiting time for the emergence of the first three SARS-CoV-2 VOC is highlighted in red. Both scenarios produce roughly the same of number of VOC lineages. However, on average, *T*_0_ is slightly longer for the *K*=6 scenario. **(B)** Evaluating the temporal clustering of the first three VOC lineages. For each simulation run, represented by a point on the graph, we measure the time that it takes for a single adaptive mutation to establish in the population and the time difference between the establishment of the first and third successful VOC lineage. The red dashed rectangle shows the region of the parameter space corresponding to the emergence of the first three SARS-CoV-2 VOCs with the cross sign (“**X**”) representing the mean value. We can see that a noticeable fraction of simulation runs for both scenarios yield a temporal clustering that matches the emergence of the first three VOCs in late 2020 (i.e., they fall inside the enclosed area). The inset shows that 35.5% and 25.9% of the runs for *K*=3 and 6 scenarios produce fewer than three successful VOC lineages by the end of the simulation period, respectively. Each run stops once the frequency of the VOC population reaches 75%. **(C)** A typical evolutionary trajectory corresponding to the *K*=6 scenario highlighted with a bold orange circle in panel (B). The graph shows the background population in gray and the *i*-mutant populations (1<*i*≤*K*) in different shades of orange from light (fewer mutations) to dark (more mutations). The dashed lines show the dynamics of all the established VOC lineages over time. Red vertical arrows show the establishment time of the first three VOCs. We can see that the single-mutant genotypes (lines in light orange) are produced via the between-host pathway from very early on in the pandemic but are at very low prevalence before the emergence of the VOCs (white dashed lines). See also supplementary figure 6.

## Discussion

The global spread of the Omicron variant of SARS-CoV-2 has given a renewed attention to the underlying evolutionary mechanisms that lead to the emergence of VOCs. Practically, we would like to know whether to expect future VOCs to arise, and if so when and whether there will be early warning signs. Answering this question is not only important for understanding the fate of the pandemic but also may have major public health implications for how to best develop strategies for controlling the spread of the disease. In this study, we provided a quantitative framework for investigating the likelihood of different evolutionary pathways that can give rise to VOCs of SARS-CoV-2. We found that VOCs are unlikely to be driven by a single adaptive change at the population level as this would require significantly high levels of overdispersion which is not supported by any existing epidemiological study on SARS-CoV-2 transmission [25]. We also showed that if multiple VOC mutations combine additively for advantage, they can only emerge on the background of a chronic infection, otherwise individual VOC mutations would reach high frequencies from the early stages of the pandemic and, therefore, would have been picked up from genomic surveillance data. If individual VOC mutations were acquired during chronic infections and had a strong advantage relative to the then-dominant genotypes, they may have still been leaked to the population at large before the emergence of VOCs. Therefore, we showed that only additive mutations with relatively small fractional contribution to VOC fitness may yield evolutionary dynamics that resembles the clustered emergence of SARS-CoV-2 VOCs in late 2020. On the other hand, we showed that cryptic circulation of a mutant lineage for sustained periods of time before producing VOCs is possible via a fitness plateau-crossing landscape. While at the between-host level such a landscape may not yield more than 2 mutations in excess of the background population over a period of 7-12 months under biologically relevant parameter combinations, many more mutations can be accumulated during a chronic infection, for example such as those found in certain immunocompromised individuals, without ever being leaked to the rest of the population. We found that the pattern of the timing of VOC emergence via the fitness plateau-crossing landscape under both the within- and between-host pathways are aligned with the timing of the clustered emergence of Alpha, Beta, and Gamma variants in late 2020.

Finally, it is important to note that in all of the within-host evolutionary pathways (i.e., Landscapes 1, 2, and 3), we found parameter combinations that can re-create the clustered emergence of the first three SARS-CoV-2 VOCs. In particular, we showed that either because of having very few mutations that are selectively beneficial at the population level (i.e., Landscape 1) or the low prevalence of intermediate mutations before the emergence of the VOCs (i.e., Landscapes 2 and 3), we would expect the phylogenetic relationship between the VOC lineages and background populations to manifest itself with a long evolutionary distance branch connected to deeper internal nodes of the tree with each VOC clade independently emerging from a unique genetic background and be subsequently replaced by another VOC clade from an entirely different background (**Figure 9**). This creates a phylogenetic relationship between VOC clades that is similar to what we observe for Alpha, Beta, and Gamma variants [4–6].

**Figure 9:**
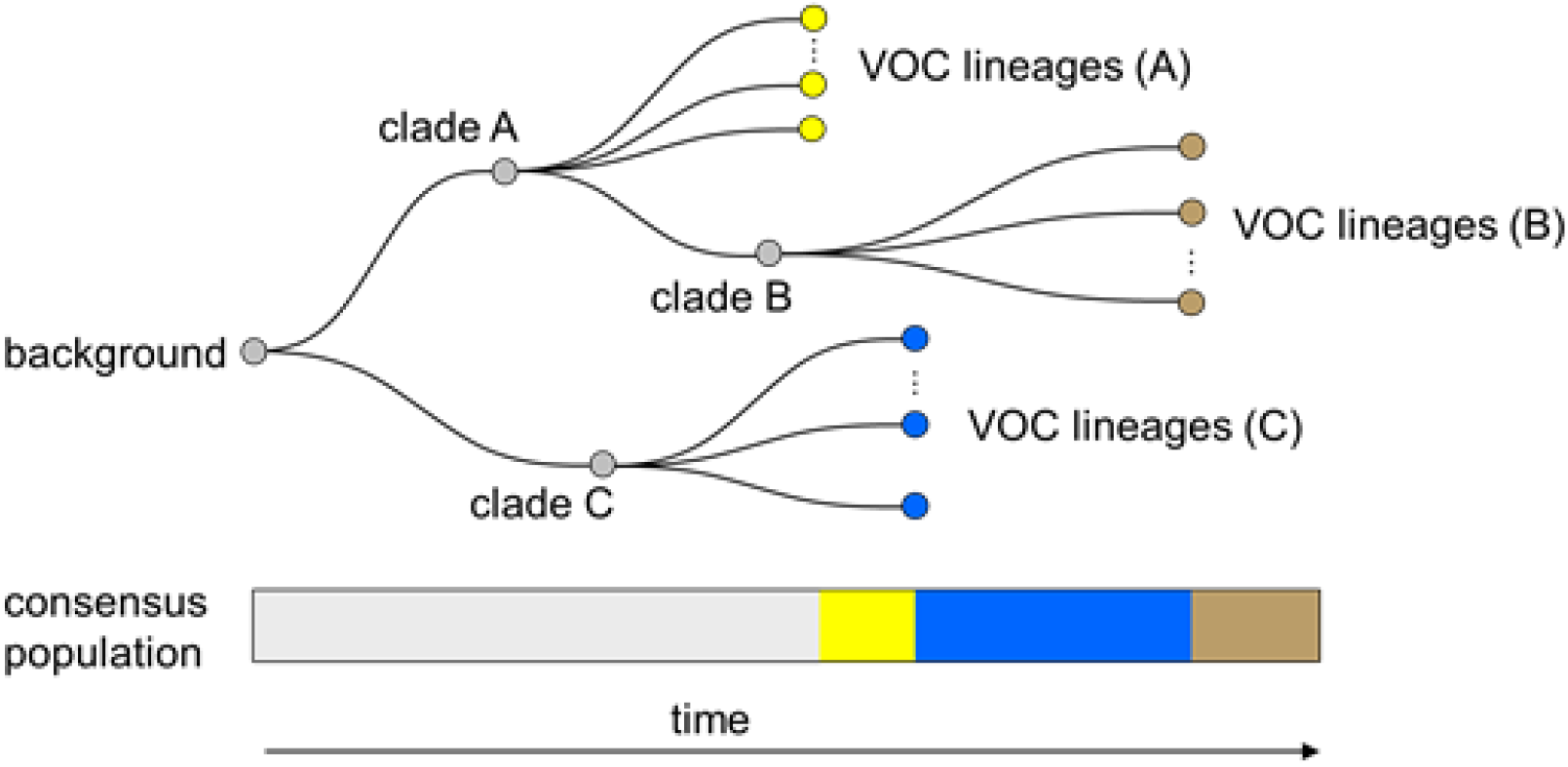
Within-host evolution reproduces the star-like genealogy of the VOCs. For a wide range of parameter combinations in Landscape 1, 2, and 3, we showed that the within-host evolutionary dynamics can become virtually uncoupled from the the evolution at the between-host level enabling each VOC lineage to arise independently on the background of a different clade which leads to a star-like tree topology.

### Cryptic transmission of VOCs in humans

Another possibility for why the VOCs were not detected until mid to late 2020 is that they may have been circulating cryptically in areas of the world with poor genomic surveillance before becoming globally dominant. While variants of SARS-CoV-2 with multiple spike mutations have been detected through travel surveillance from passengers travelling from areas with little to no genomic surveillance [27–28], if they were highly transmissible variants and had a potential to become a VOC, given the interconnectivity of the human interactions, it should not take very long before they become globally dominant. Therefore, as we showed in our analysis of Landscape 1, this scenario of VOC emergence seems to be only possible under significant levels of overdispersion such that it prohibits the selectively beneficial mutations from immediately taking off globally relative to other variants.

### Possibility of reverse zoonosis

A somewhat similar idea to the cryptic transmission of variants in human populations is the possibility of a lineage (or multiple lineages) of SARS-CoV-2 jumping from humans to other mammals such as white-tailed deer, mink, hamster, and mouse where they circulate and evolve without being detected for a relatively long period before they jump back to the human population [29–32]. In particular, some recent studies have reported the detection of multiple spillovers of SARS-CoV-2 from humans and onward transmission in deer population with highly divergent genomes being detected in deer population with potential deer-to-human transmission [33–34]. However, the genomic composition of these divergent genomes in deer population are different from the VOCs with a much lower ratio of non-synonymous to synonymous changes which suggests they may be following a completely different evolutionary path. Nevertheless, these studies indicate that it is possible for a highly divergent set of genomes to evolve in another species with a potential for deer-to-human transmission without ever being detected. Mink and hamster sequences offer some of the more compelling examples of transmission from humans to a non-human species and back, supported by phylogenetic evidence [30–31]. None of the currently identified sequences from animals appear as sister taxa to any of the circulating VOCs. While we cannot rule out evolution in an animal reservoir, one might expect the contribution of animals to human transmission chains to be dwarfed by the amount of human-to-human transmission currently happening.

### Role of recombination

Recombination can bring together mutations from different backgrounds, potentially expediting the rate of adaptation by creating viable and more pathogenic hybrid new variants of a pathogen. Coronaviruses are also known to recombine with one another during mixed infections [35–36]. While during the early stages of the pandemic SARS-CoV-2 sequences typically differed by only a handful of mutations from each other thereby making the effects of recombination indistinguishable from those of recurrent mutation [37], as more viral genetic diversity built-up in the population, the generation and transmission of interlineage recombinants of SARS-CoV-2 in humans were reported in multiple studies [38–39]. Even though there is currently no definitive evidence for recombination being involved in the emergence of VOCs, including the Omicron variant of SARS-CoV-2 [40], we would expect the role of recombination to be more pronounced as the virus continues accumulate more genetic diversity by sustained circulation around the world.

### Shifting landscape

We have assumed a static fitness landscape prior to the emergence of the first three VOCs; here we consider the plausibility of that assumption. During the first year of the pandemic, a novel virus was spreading in an immunologically naïve population [12]. As more individuals became infected and developed natural immunity, it is possible that the fitness landscape for the virus shifted as selection for immune escape increased [41]. However, by the time the first three VOCs emerged in late 2020, the majority of the world’s population were still susceptible to the disease and may not have even been exposed to it. Therefore, it is unlikely that the build-up of natural immunity alone was the reason behind their increased selective advantage. In contrast, the global dominance of Omicron in late 2021 was largely due to its immune escape properties relative to previous variants of SARS-CoV-2 [42] and, therefore, its emergence was likely the result of a changing viral fitness landscape.

### Future VOCs

Emergence of new VOCs with increased transmissibility, immune evasion properties, and virulence poses a great challenge to managing SARS-CoV-2. Based on our findings, one of the major implications of chronic infections being the main source of generating VOCs is that finding and treating chronic infections should be a top priority, not just for the benefit of chronically ill patients but also from a public health standpoint. One of the main challenges with assessing the likelihood of VOC emergence during chronic infections would be to quantify the prevalence of immunosuppressed individuals within a population and determine which forms of immunosuppression are associated with chronic infections.

Several studies have now shown evidence of recurrent SARS-CoV-2 mutations in immunocompromised patients [15–17], with some suggesting the detection of a variant-like lineage which arose from a chronic infection that spilled over into a local population [43]. Another major implication of our work is that we can now quantitatively explain the possibility of such events and find the expected time that it takes for a new VOC to emerge from a within-host evolutionary pathway that involves any number of mutations. We showed that a typical within-host plateau-crossing or additive mutation pathway involving 3-6 mutations requires a within-host fixation rate of *μ*_C_~0.1-0.3 per generation which corresponds to a period of 50-300 days since the start of a chronic infection. The timing of such an event aligns with the time frame over which some of the major mutations involved in VOCs have been observed in patients with chronic infections [15–17]. This also implies that if a VOC emerges from the within-host evolutionary pathway, it is more likely to reflect the genetic diversity of the virus population from several months ago. It can explain why, for instance, the Omicron variants were not descendents of Delta, which was the most prevalent variant at the time of emergence of Omicron. It also suggests that while the next VOC could emerge from the prevalent Omicron background, it could also come from, e.g., a chronic infection with Delta that started prior to the Omicron wave. Some of the key remaining questions involve how much more of the fitness landscape the virus will be able to explore as more chronic cases accumulate and existing chronic cases last longer. For instance, if it has already crossed a 6-mutant fitness-plateau, how much longer would it take to explore 7-mutant fitness plateaus?

## Methods

### Between-host model of VOC emergence

#### Effective population size

We approximate the between-host evolution of SARS-CoV-2 as a haploid population of size *N*(*t*) which is equal to the number of daily infectious individuals with SARS-CoV-2 worldwide. Since the number of confirmed cases is often a significant underestimation of the true number of infections (e.g., see: [44–45]), we use the number of daily confirmed deaths [46] to back-calculate the number of infectious individuals, *N*(*t*), from the global median infection fatality rate (IFR) of COVID-19 [47]. We note this approach is still subject to several potential sources of bias including variation in IFR over time (e.g. due to various pharmaceutical interventions) and across different demographics [48]. Using confirmed COVID-19-related deaths may still underestimate the true number of deaths associated with the disease due to under-reporting of deaths particularly in areas of the world where there is limited testing from suspect cases [49]. Nevertheless, by allowing for a wide variation in global IFR (from 0.2% to 1.5%), we can capture most of the uncertainty in the number of infectious individuals worldwide. We also note that for the timespan of interest in our work (i.e., start of the pandemic until the emergence of the first three VOCs), the impact of pharmaceutical interventions such as vaccination on lowering the global IFR is likely to have been negligible given that vaccination campaigns mostly started in 2021. The confirmed global deaths started being reported from 2020-01-23. Assuming a 20-day delay from the onset of symptoms to death [50], we set 2020-01-03 as the first timepoint in the simulation.

#### Advantage of mutants

The selective coefficient of a mutant individual depends on its number of mutations and the fitness landscape (see **Figure 2A**). For instance, in the case of an additive fitness landscape of size *K*=3, the fitness advantage of the single-, double-, and triple-mutants are *s*/3, 2*s*/3, and *s* relative to the wild-type population, respectively. During one generation, the frequency *f_i_* of individuals with genotype *i* and selective advantage *s_i_* relative to the wild-type increases by a factor (1 + *s_i_*), along with further adjustments to their frequency due to mutations from/to other genotypes. Upon normalisation (∑*_i_ f_i_* = 1), these frequencies are used for the Dirichlet-multinomial sampling step. After the sampling step, the numbers of cases are converted to frequencies for sampling in the next generation.

#### Epidemic spread

Due to a high degree of individual-level variation in the transmission of SARS-CoV-2 (i.e., overdispersion) [25–51], we use a Dirichlet-multinomial (instead of a multinomial) distribution to assign offspring in generation *t*+1 to parents from generation *t*. The Dirichlet-multinomial is parametrized by *N*(*t*+1) (the number of offspring to draw for the next generation) and *A* 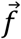, the weights of the different genotypes, where 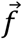 is the normalized vector giving their frequencies in the current generation. The scalar *A* controls the amount of dispersion, with smaller *A* corresponding to increased demographic noise. To match it to observations, we note that under the Dirichlet-multinomial model, the number of secondary cases produced by an infection approximately follows a negative binomial distribution with mean *R_t_* = *N(t+1) / N(t)*. In terms of the Dirichlet multinomial parameters, the variance of this negative binomial is 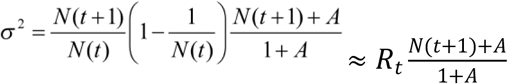. This should match the variance in the number of secondary cases written in terms of the dispersion parameter *k*, *σ^2^* = *R_t_(1 + R_t_/k)*. Equating these two expressions gives 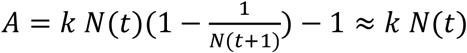.

#### Mutation rate

Assuming a constant generation time 5.2 days for all variants of SARS-CoV-2 over time [52], we use the phylogenetically estimated substitution rate per site per year [53] to calculate the mutation rate per site per generation time, *μ*. We also note that generation time may vary over time depending on the behavioural changes in the population and emergence of variants, that is why we allow for some variation in the mutation rate parameter in our model (0.87 - 2.0) x 10^-5^ based on phylogenetic estimates. We assume each site has two states: wild-type and mutant. Therefore, for a group of *K* sites, there are 2*^K^* genotypes.

#### Inferring the selective advantage of VOCs

Finally, the selective advantage *s* of the VOCs is determined by fitting an exponential function, *f*(*t*), of the form, *f*(*t*)=*ae^st^*, to the proportion of Alpha, Beta, and Gamma variants sampled in the country where they were first detected (i.e., UK, South Africa, and Brazil) using the NonlinearModelFit function in Mathematica 11.0 (cite Mathematica). We find that the selective advantage *s* for Alpha, Beta, and Gamma are 0.37 (95% confidence interval: 0.33 - 0.41), 0.74 (95% confidence interval: 0.65 - 0.83), and 0.84 (95% confidence interval: 0.58 - 1.08), respectively (see **Supplementary Figure 7**). The confidence interval is obtained by multiplying the standard error by the value of Student’s t for the given confidence level and degrees of freedom. Given the uncertainty in our estimates due to noise in the observations, potential sampling bias, and spatio-temporal heterogeneities, we make the assumption that the value of *s* is roughly the same for the different VOCs and use the same estimate for all three trajectories (**Table 1**).

### Within-host model of VOC emergence

Each VOC mutation is fixed within the host at rate *μC* such that the fixation time is an exponentially distributed number with mean 1/*μ_C_*. Each mutation may then spread to the rest of the population with a probability that is proportional to its fitness as determined by the Dirichlet-multinomial sampling. At any time during the pandemic, a chronic infection can be seeded by other infectious individuals within the population, *N*(*t*), with a probability *P_f_*. Therefore, at every generation, the number of chronic infections is given by a binomial distribution with success probability *P_f_*. Once a chronic infection emerges, it remains in the population for the remainder of the simulation period.

### Simulation setup

For both within-host and between-host models of VOC emergence, we run each evolutionary scenario for a given combination of model parameters 1,000 times. We then measure total number of established VOC lineages, *M*, the time that it takes for the establishment of the first VOC, *T*_0_, and the time between the establishment of the *i*^th^ and (*i*+1)^th^ VOC, *T*_i:(i+1)_, for the first 6 established VOC lineages in each scenario. An *established* VOC lineage is defined as a lucky lineage with selective advantage *s* that survives drift upon reaching a size 1/*s*. Similarly, the establishment time of a VOC lineage is defined as the time that it takes for that lineage to reach size 1/*s*. Each run stops once the frequency of the VOC population reaches 75%.

## Code and data availability

All software code and analysis scripts to reproduce the figures are available at https://github.com/weissmanlab/Valley_Crossing.

## Supplementary figures

**Supplementary figure 1:**
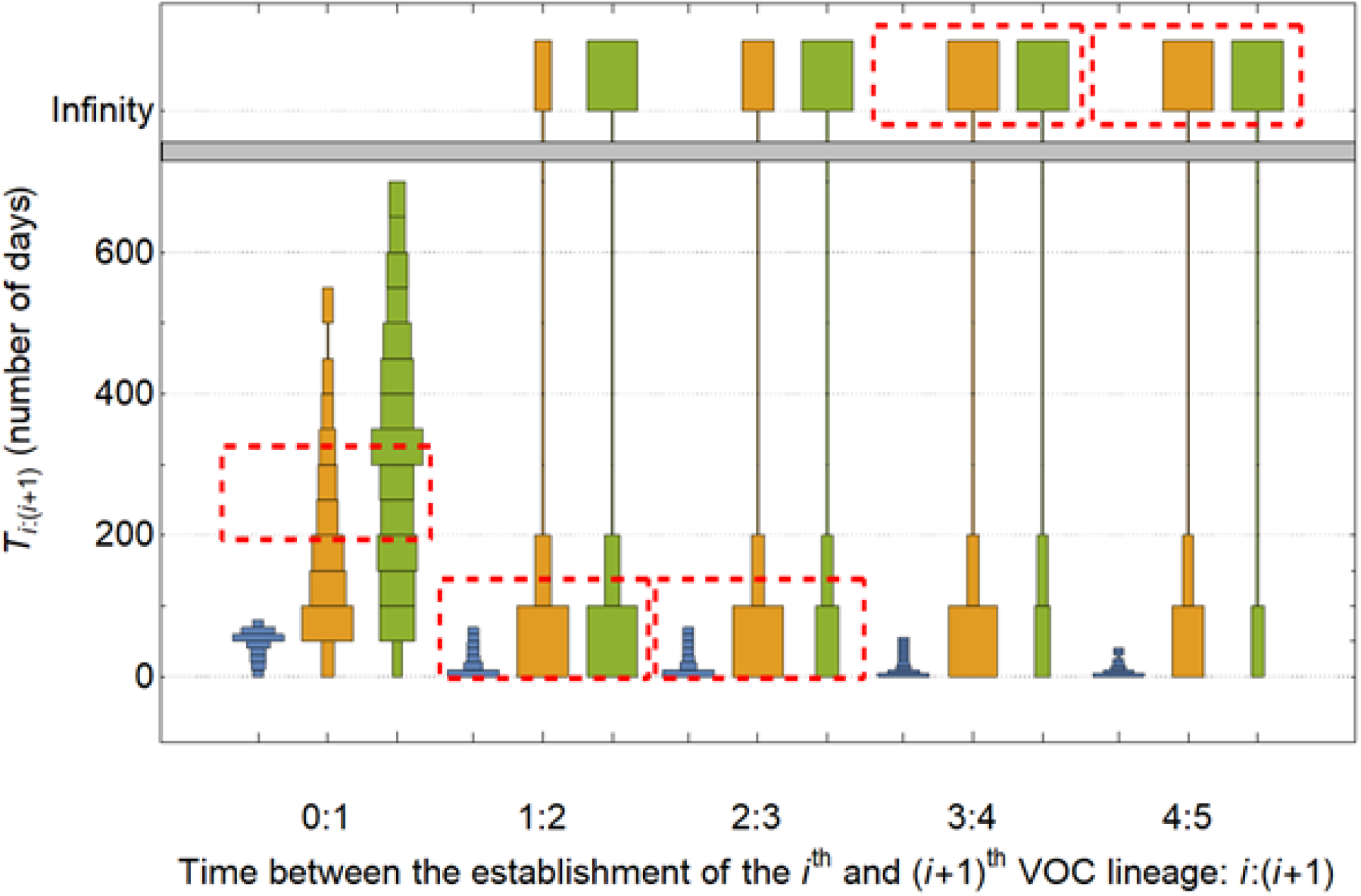
Distribution of waiting times for the establishment of consecutive pairs of VOC lineages via the between-host pathway assuming a fitness landscape with a single adaptive mutation. The distribution of times that it takes between the production of the *i*^th^ and (*i*+1)^th^ lineage, *T*_i:(i+1)_, for the first 5 established VOC lineages described in Figure 3. *T*_0:1_ is the waiting time for the production for the establishment of the first VOC lineage (equivalent to *T*_0_).

**Supplementary figure 2:**
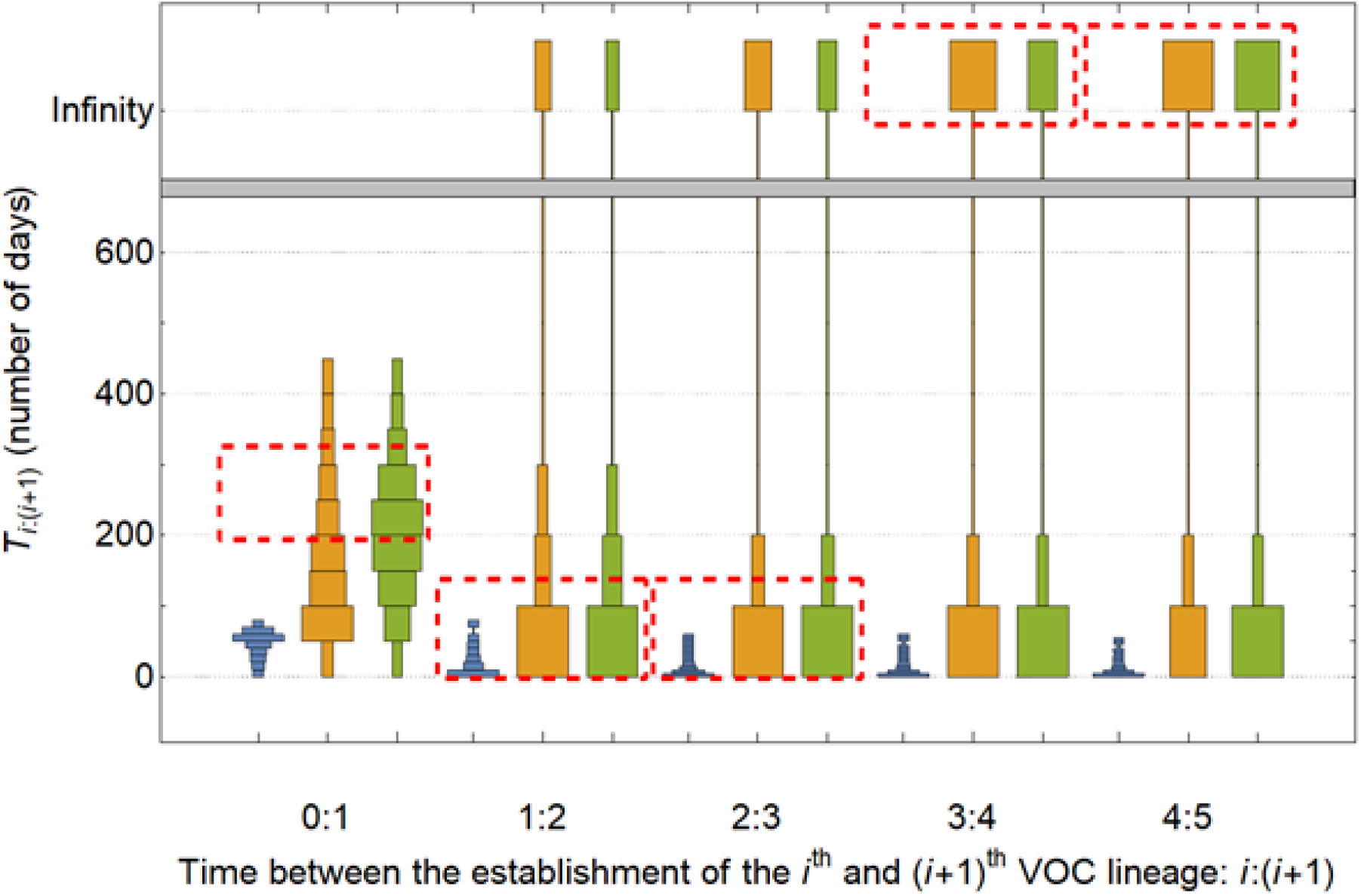
Distribution of waiting times for the establishment of consecutive pairs of VOC lineages via the within-host pathway assuming a fitness landscape with a single adaptive mutation. The distribution of times that it takes between the production of the *i*^th^ and (*i*+1)^th^ lineage, *T*_i:(i+1)_, for the first 5 established VOC lineages described in Figure 4. *T*_0:1_ is the waiting time for the production for the establishment of the first VOC lineage (equivalent to *T*_0_).

**Supplementary figure 3:**
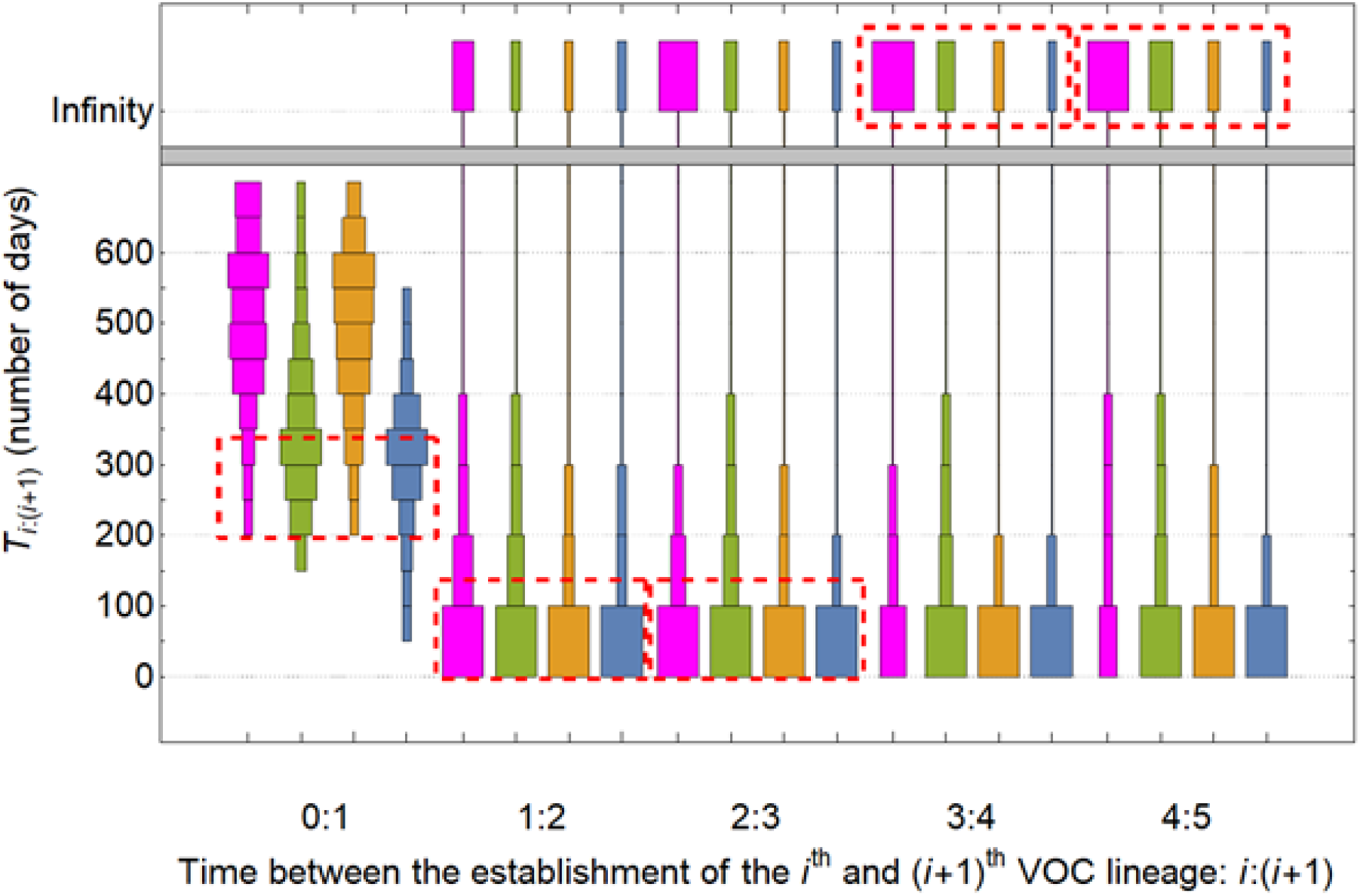
Distribution of waiting times for the establishment of consecutive pairs of VOC lineages via the between-host pathway assuming an additive fitness landscape. The distribution of times that it takes between the production of the *i*^th^ and (*i*+1)^th^ lineage, *T*_i:(i+1)_, for the first 5 established VOC lineages described in Figure 5. *T*_0:1_ is the waiting time for the production for the establishment of the first VOC lineage (equivalent to *T*_0_).

**Supplementary figure 4:**
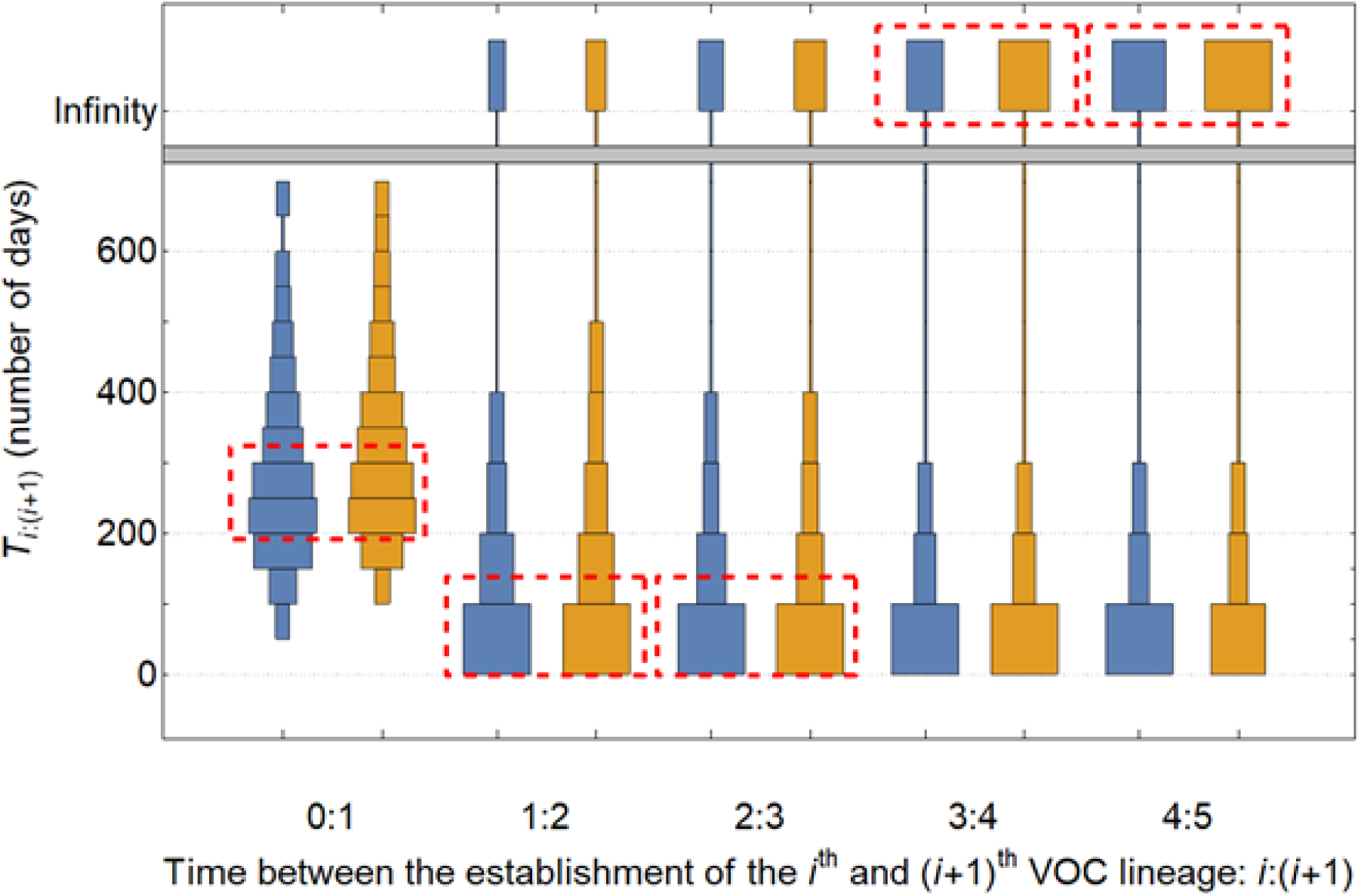
Distribution of waiting times for the establishment of consecutive pairs of VOC lineages via the within-host pathway assuming an additive fitness landscape. The distribution of times that it takes between the production of the *i*^th^ and (*i*+1)^th^ lineage, *T*_i:(i+1)_, for the first 5 established VOC lineages described in Figure 6. *T*_0:1_ is the waiting time for the production for the establishment of the first VOC lineage (equivalent to *T*_0_).

**Supplementary figure 5:**
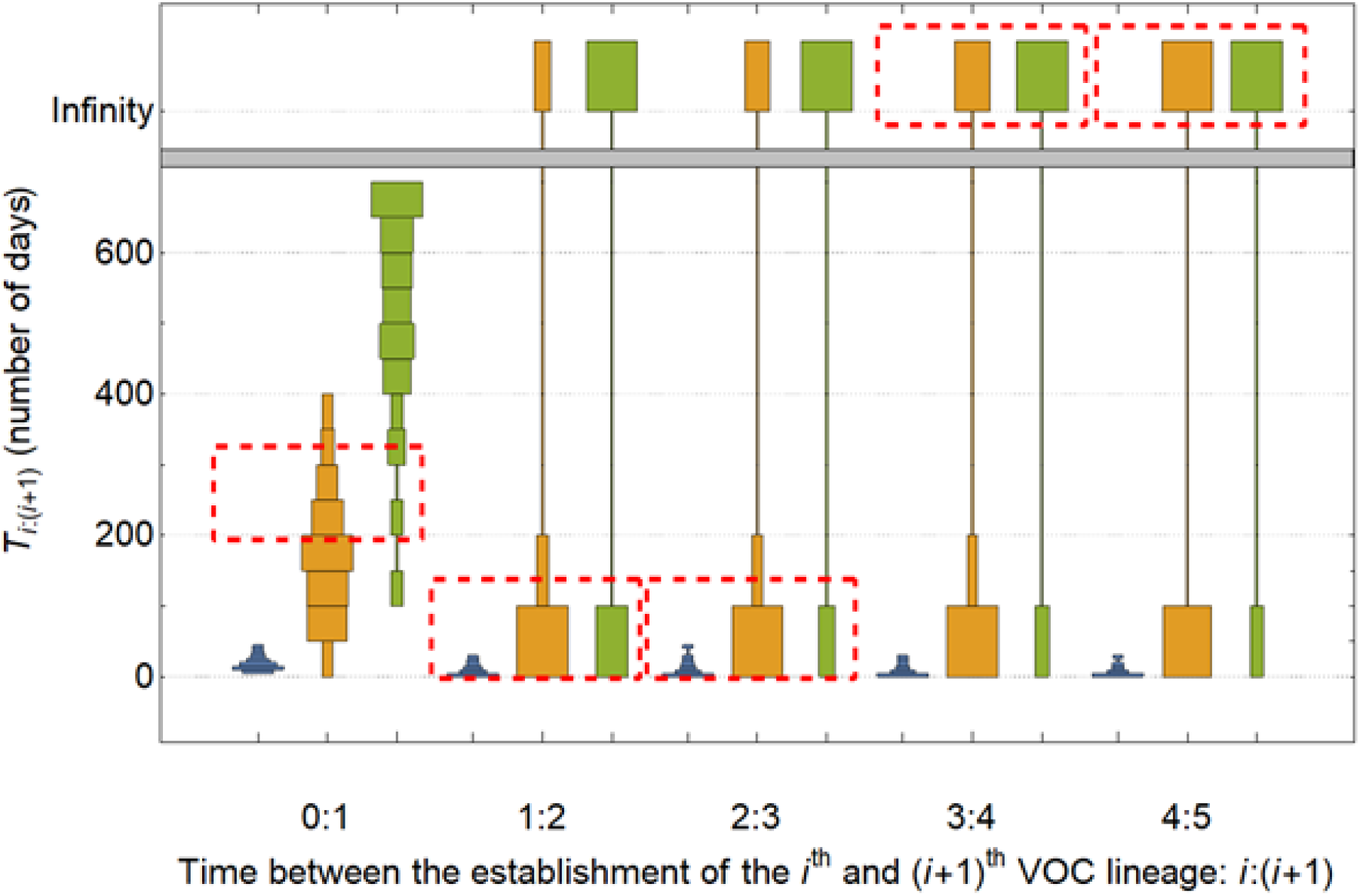
Distribution of waiting times for the establishment of consecutive pairs of VOC lineages via the between-host pathway assuming a fitness plateau landscape. The distribution of times that it takes between the production of the *i*^th^ and (*i*+1)^th^ lineage, *T*_i:(i+1)_, for the first 5 established VOC lineages described in Figure 7. *T*_0:1_ is the waiting time for the production for the establishment of the first VOC lineage (equivalent to *T*_0_).

**Supplementary figure 6:**
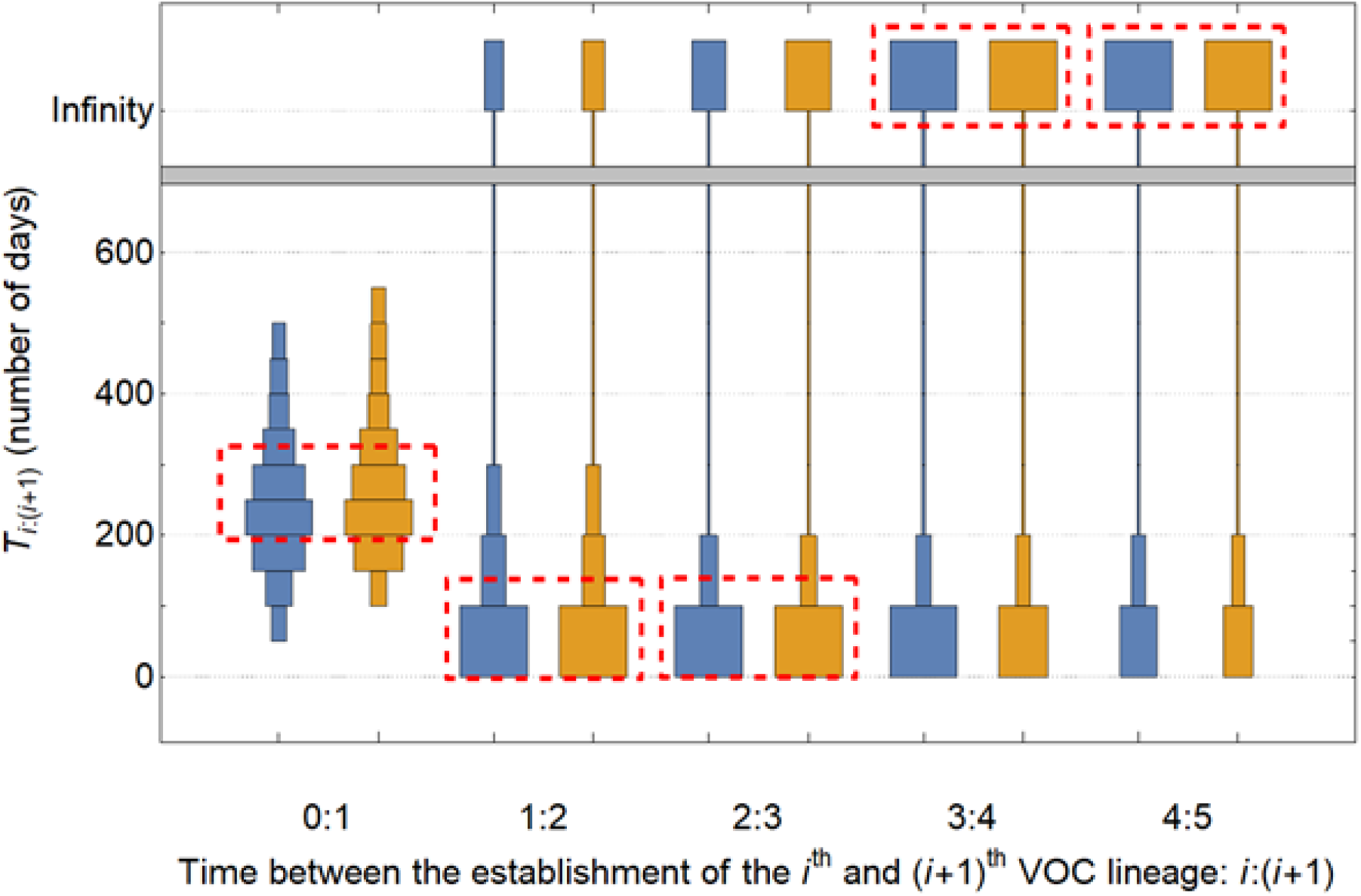
Distribution of waiting times for the establishment of consecutive pairs of VOC lineages via the within-host pathway assuming a fitness plateau landscape. The distribution of times that it takes between the production of the *i*^th^ and (*i*+1)^th^ lineage, *T*_i:(i+1)_, for the first 5 established VOC lineages described in Figure 8. *T*_0:1_ is the waiting time for the production for the establishment of the first VOC lineage (equivalent to *T*_0_).

**Supplementary figure 7:**
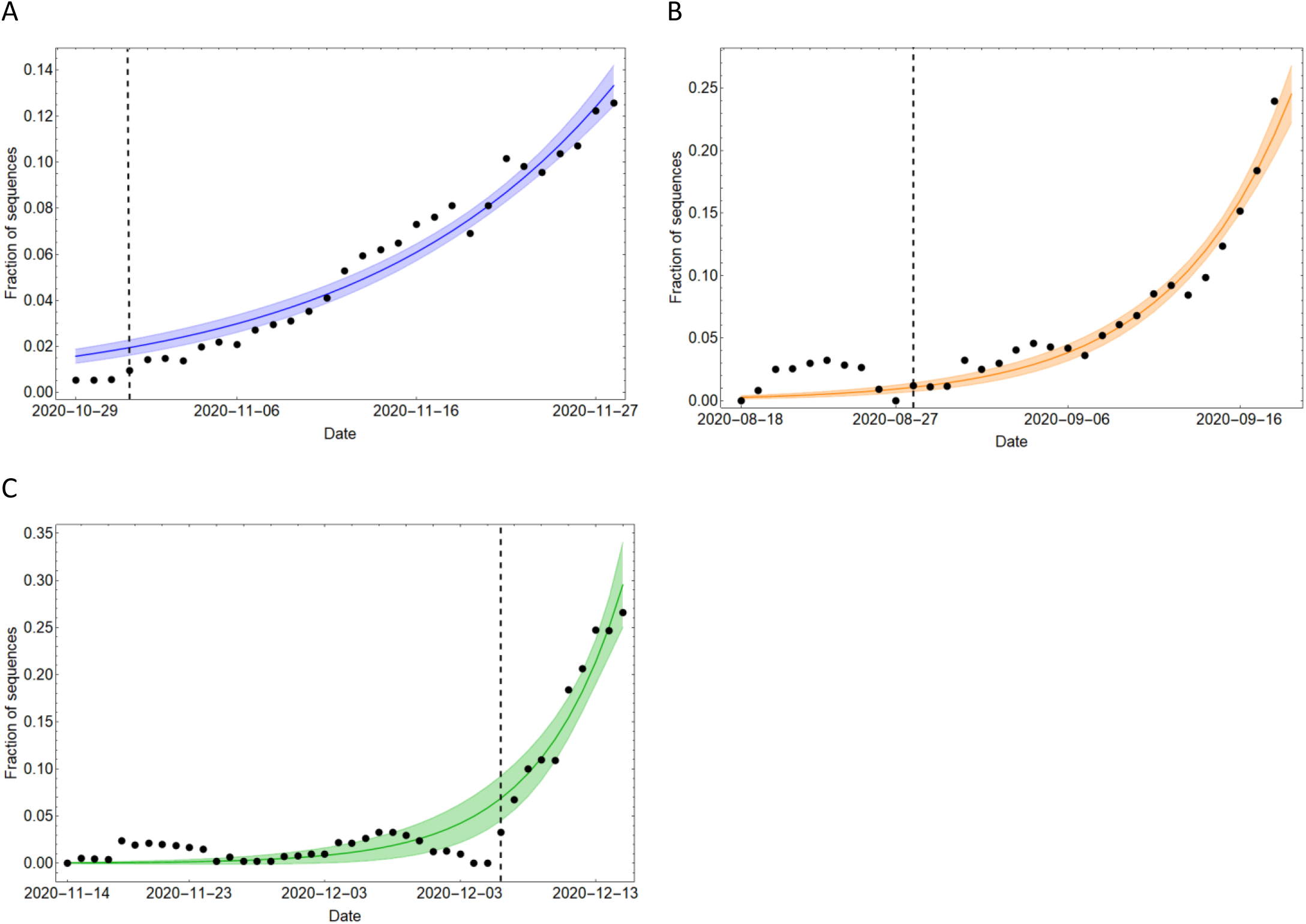
Exponential model fits to the frequency of individual SARS-CoV-2 VOC sequences sampled in its country of origin. **(A-C)** Fitting an exponential function of the form, *f*(*t*)=*ae^bt^*, to the frequency of Alpha, Beta, and Gamma sequences sampled in the UK, South Africa, and Brazil, respectively. Vertical dashed line shows the starting timepoint used for the fitting. The shaded area shows the mean prediction bands.

